# Dephosphorylation of YES kinase-mediated co-chaperone DNAJB6b phosphorylation attenuates tau protein aggregation

**DOI:** 10.1101/2025.11.14.688390

**Authors:** Meng-Chieh Lin, Huan-Yun Chen, Te-Sheng Lin, Sue-Jane Lin, Pei-Jie Lin, Chia-Yu Liao, Nei-Li Chan, Chin-Hsien Lin, Shu-Chun Teng

## Abstract

Alzheimer’s disease (AD) is a neurodegenerative disorder characterized by a gradual deterioration of memory. Here, we examine the biological consequences of phosphorylation-mediated chaperone activity in AD-associated tau aggregates. Increased phosphorylation of DNAJB6b at Y53 is observed in the brain lysates of AD patients. Our research found that an activated Src family kinase, YES, phosphorylates Y53 within the J-domain of DNAJB6b and enhances the binding between DNAJB6b and Hsp70. The strengthened association between DNAJB6b and Hsp70 may lead to the accumulation of tau aggregates. These findings suggest that the up-regulation of YES kinase modifies DNAJB6b, and that the resulting alteration of DNAJB6b-dependent tau disaggregation may contribute to an increased risk of developing AD. Additionally, phosphor-DNAJB6b at Y53 may serve as a biomarker for the prognosis and diagnosis of AD.

## Introduction

AD, a common neurodegenerative disease, is characterized by cognitive impairment and memory decline [1]. The pathological hallmarks of aging-related neurodegenerative disease [2] are the loss of proteostasis and the accumulation of toxic aggregates. Specific abnormal AD structures are the deposits of intracellular hyperphosphorylated tau protein aggregates and extracellular amyloid-β plaque in the hippocampus [3]. Tau protein, belonging to the microtubule-associated protein (MAP) family, normally facilitates the assembly of microtubules and stabilizes the microtubule dynamics. However, abnormal folding of tau protein can increase its propensity for self-aggregation and lead to the formation of intracellular neurofibrillary tangles (NFTs), which are the primary pathological hallmarks of AD and cause cognitive decline. Aggregated insoluble tau protein in cells impairs a broad range of cellular processes, including axonal transportation [4], per- and pro-synapse transmission [5], and mitochondrial function [6], which collectively lead to profound damage to neuronal health. Additionally, misfolded tau protein ‘spreads’ its toxicity to other native tau molecules and induces aberrant conformation change [7]. The pathological progression of AD can be classified into six stages, proposed by Braak [8], based on the distribution pattern of neurofibrillary change. The progression begins with the formation of early-stage intraneuronal tau aggregates in the transentorhinal cortex and culminates in advanced isocortical degeneration driven by high-density neurofibrillary tangles. This hierarchical spread of tau pathology correlates with subtle memory impairments, sequential cognitive decline, and advancing to severe cortical dysfunction.

The heat shock protein 70 (Hsp70) system is constitutively active and critical for maintaining cellular proteostasis by aiding in the correct folding of proteins or directing misfolded or aggregation-prone proteins toward degradation [9, 10]. To activate a Hsp70 reaction machinery cycle, it requires the active participation of Hsp70s, co-chaperone J-domain proteins (JDP), nucleotide exchange factors (NEF), and ATP-ADP exchange [10, 11]. Human Hsp70 comprises 13 compartment-specific isoforms that fulfill organelle-specific functions. In the cytoplasm, it includes stress-inducible HSPA1A/HSP70 and constitutively expressed HSPA8/HSC70 [12, 13]. The Hsp70 exhibits intrinsically low ATPase activity in its unliganded state, requiring JDPs for functional activation [14]. JDPs perform dual roles: delivering substrates to Hsp70’s substrate-binding domain and allosterically stimulating ATP hydrolysis through conserved J-domain interaction. This ATPase-mediated activation induces structural transitions in Hsp70, generating mechanical forces for substrate folding. The coordinated JDP-mediated substrate refolding and NEF-driven nucleotide exchange establish an iterative folding mechanism, demonstrating an essential partnership in proteostasis. This ATP-dependent cycle highlights how JDPs critically regulate substrate and ATPase activation, ensuring efficient protein quality control.

Over 50 JDPs found in humans can be classified into three subfamilies: DNAJA, DNAJB, and DNAJC [15]. Many findings implicate that the DNAJB subfamily is associated with delivering distinct types of misfolded clients [16]. DNAJB1, for example, is proven to slow the spreading of α-synuclein aggregation associated with Parkinson’s disease [17, 18]. Misfolding-prone A25T mutant transthyretin (TTR) and amyloid precursor protein (APP) can be bound to secreted DNAJB11 to prevent extracellular aggregation and proteotoxicity during ER stress [19]. Polyglutamine aggregation from Huntington’s disease can be inhibited by DNAJB6, both *in vitro* and *in vivo* [20]. Previous studies have also shown that DNAJB6 expression is associated with the aggregate load of tau in neuroblastoma cells [21]. Reduced expression of DNAJB6b is correspondingly observed with the increase of tau aggregation in an age-dependent manner in an rTg4510 mouse model [21]. DNAJB6 is ubiquitously expressed across various tissues, including the brain and muscle. It has two alternative splicing isoforms, nuclear-localized a form (36 kDa) and cytoplasmic-localized b form (27 kDa) [22]. Structurally, like other DNAJB subfamilies, DNAJB6 contains an N-terminal J-domain followed by a glycine/phenylalanine (G/F)-rich, a serine/threonine (S/T)-rich, and a functionally unclear C-terminal domain [23]. J-domain is a signature domain of JDPs with a His-Pro-Asp (HPD) motif required for direct interaction with Hsp70 [24]. G/F-rich and S/T-rich regions are thought to participate in client recognition and aggregate suppression, respectively [20, 25]. Despite the importance of the JDP-Hsp70 system in maintaining cellular proteostasis networks, current understanding of how posttranslational modifications (PTMs) of JDP regulate the chaperone system remains limited.

Over 90% of ADs are not caused by genetic mutations [26], suggesting that environmental, post-transcriptional, and post-translational factors during the aging process may be the key regulators. Hundreds of PTMs on chaperones/co-chaperones, known as the chaperone codes, are thought to alter chaperone activity and protein interactions [27]; however, our understanding of the functional roles of the chaperone codes is still limited [18, 28, 29]. Combined with the existing knowledge of the specific association between DNAJB6b and tau protein, here we explore the influence of phosphorylation on regulating the co-chaperone activity of DNAJB6b.

## Results

### Dephosphorylation of DNAJB6b on tyrosine 53 enhances its ability to prevent tau aggregation in SH-SY5Y neuroblastoma cells

Previous studies have shown that co-chaperone DNAJB6b is associated with the aggregate load of tau in neuroblastoma cells [21]. To determine the functional role of the chaperone code on DNAJB6b, we considered candidates based on available data from PhosphositePlus® [30] (see **Table S1**). As the J-domain of DNAJB6b is essential for preventing tau aggregation, phosphorylation of tyrosine 53 (Y53), as determined by the highest number of references on the J domain using proteomic mass spectrometry, was then chosen for further investigation. To evaluate the effect of the phosphomimetic substitutions in cells, the Y53 site on DNAJB6b is replaced by phenylalanine (F) (non-phosphorylatable variants) or glutamic acid (E) (pseudo-phosphorylated mutants). The filter trap assay was first used to detect and quantify protein aggregation levels. Human neuroblastoma SH-SY5Y cells overexpressing the disease-associated EGFP-tagged tau P301L exhibited a significant tau aggregation level. However, under the co-overexpression of V5-tagged DNAJB6b wild-type, tau aggregates were reduced (**Figure 1A-B**), consistent with previous observations. To further determine whether the phosphorylation of DNAJB6b Y53 affects its ability to reduce tau aggregation, we co-overexpressed EGFP-tau P301L with either non-phosphorylatable V5-DNAJB6b Y53F or pseudo-phosphorylated V5-DNAJB6b Y53E in SH-SY5Y cells. A decrease in aggregated tau was observed in the V5-DNAJB6b Y53F mutant group. In contrast, the V5-DNAJB6b Y53E mutant displayed no significant difference compared to that in the V5-DNAJB6b wild-type group (**Figure 1C-D**), suggesting that dephosphorylation of Y53 may strengthen the function of DNAJB6b and associate with fewer tau aggregates.

**Figure 1.**
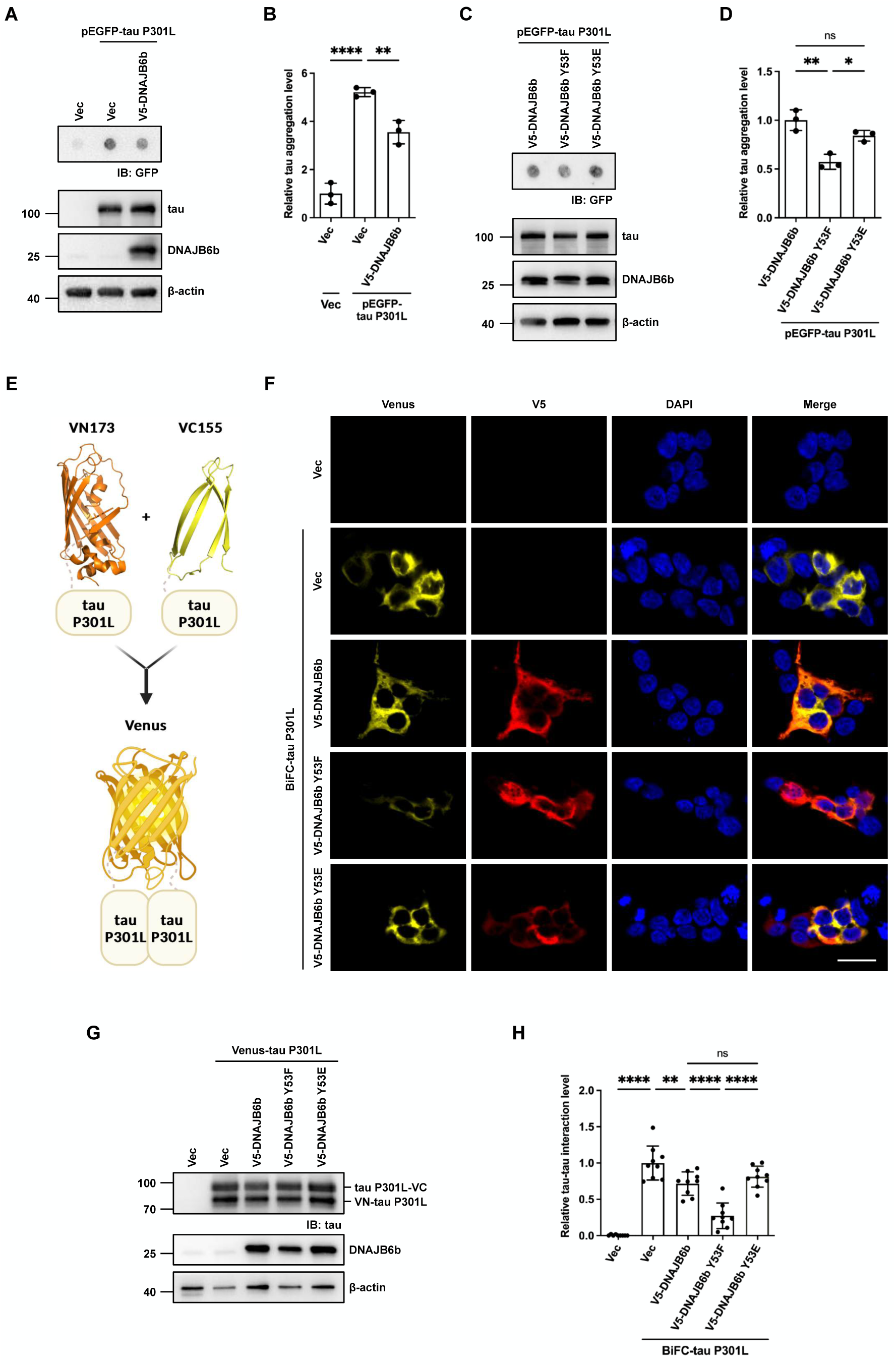
Dephosphorylation of DNAJB6b enhances its ability to prevent tau aggregation. **(A)** The filter trap assay for analysis and quantification of aggregated tau proteins. Retained proteins were immunoblotted using an anti-GFP antibody to detect tagged tau. **(B)** β-actin served as a loading control for quantification of aggregated tau proteins in **(A)** (*n*=3). **(C)** SH-SY5Y cells co-transfected with EGFP-tau P301L and different combinations of the V5-DNAJB6b Y53 constructs for 48 hours using the same filter trap assay method as in **(A)**. **(D)** β-actin was used as a loading control for quantification of aggregated tau proteins in (**C)** (*n*=3). **(E)** Schematic representation of BiFC maturation for visualization of tau aggregation. Non-fluorescent N-terminal (VN173) and C-terminal (VC155) compartments of Venus protein fused to full-length tau P301L. Aggregation of the tau protein leads to fluorescence emission. **(F)** Representative fluorescent images of the BiFC assay. SH-SY5Y cells were transfected with VN-tau P301L and tau P301L-VC (BiFC-tau P301L), along with different V5-DNAJB6b Y53 constructs. The fluorescence emitted by Venus complementation was recorded 48 hours after transfection. Cells were immunostained with V5 antibody, and nuclei were counterstained with DAPI. Representative images were taken at 40x magnification. Scale bar: 20 μm. **(G)** Immunoblotting with anti-tau and anti-DNAJB6 antibodies indicated the expression level of BiFC-tau P301L and DNAJB6b in cells, respectively. **(H)** Quantification of BiFC-fluorescence of **(F)** (*n*=3). The percentage of fluorescent (BiFC^+^) SH-SY5Y cells to the total number of cells (DAPI^+^) was quantified in at least three random-field images taken at 10x magnification. Data in **(B, D, H)** were represented as mean ± SD, and *p*-values were calculated via one-way ANOVA followed by Dunnett’s test. The individual replicate data in (**B**), (**D**), and (**H**) are listed in **Table S10**. (ns, not significant; * *p* < 0.05, ** *p* < 0.01, **** *p* < 0.0001)

To confirm whether the dephosphorylation of DNAJB6b Y53 enhances co-chaperone function to prevent tau aggregation *in vivo*, a tau bimolecular fluorescence complementation (BiFC) assay was performed to evaluate the tau-tau interaction level [31, 32] (**Figure 1E-H**). Non-fluorescent N-terminal (1-172 a.a., VN173) and C-terminal (155-238 a.a., VC155) compartments of Venus protein fused to full-length tau P301L were co-overexpressed equally with V5-DNAJB6b wild-type, Y53F, or Y53E in SH-SY5Y cells (**Figure 1G**). Venus fluorescence protein complex can be formed at close distance (less than 10 nm) as tau assembles in cells (tau-BiFC signal) to visualize protein-protein interactions (**Figure 1E**). As expected, overexpression of the V5-DNAJB6b wild-type showed a decrease in the tau-BiFC signal compared to the Venus-tau P301L-only group (**Figure 1F**). Additionally, the V5-DNAJB6b Y53F-overexpressed group exhibited a significantly lower tau-BiFC positive signal than both the V5-DNAJB6b wild-type and Y53E mutants, indicating a reduced level of tau-tau interaction (**Figure 1H**). Conversely, a similar tau-BiFC positive signal was detected in the overexpression of V5-DNAJB6b wild-type and Y53E. Together, these results suggest that dephosphorylation of DNAJB6b Y53 can enhance the co-chaperone activity of DNAJB6b in decreasing the tau-tau interaction level.

### DNAJB6b Y53 phosphorylation is correlated with Alzheimer’s disease brain

Next, we examined whether phosphorylation of DNAJB6b Y53 exists in AD brains. Phosphor-Y53 antibodies were generated by immunizing a BALB/c mouse with the synthesized DNAJB6b phosphor peptide (AEApYEVLSDAKKC) and tested to verify the phosphorylation level (see **Figure S1A-B**). Western blot analysis was performed in human AD brain lysates to evaluate the phosphorylation level of DNAJB6b Y53 (**Figure 2A**). The phosphorylation of DNAJB6b Y53 was significantly increased in AD brains compared with non-AD brains (**Figure 2B**). The phosphorylation-dependent anti-tau antibody AT8 is used to detect phosphorylation of serine 202 and threonine 205 in tau from normal and AD brains. In **Figure 2C**, the quantification of the immunoblots revealed that AD brains had a higher phosphorylation level of tau, compared with non-AD brains. Therefore, DNAJB6b Y53 phosphorylation is correlated with the accumulation of tau aggregates in AD brains.

**Figure 2.**
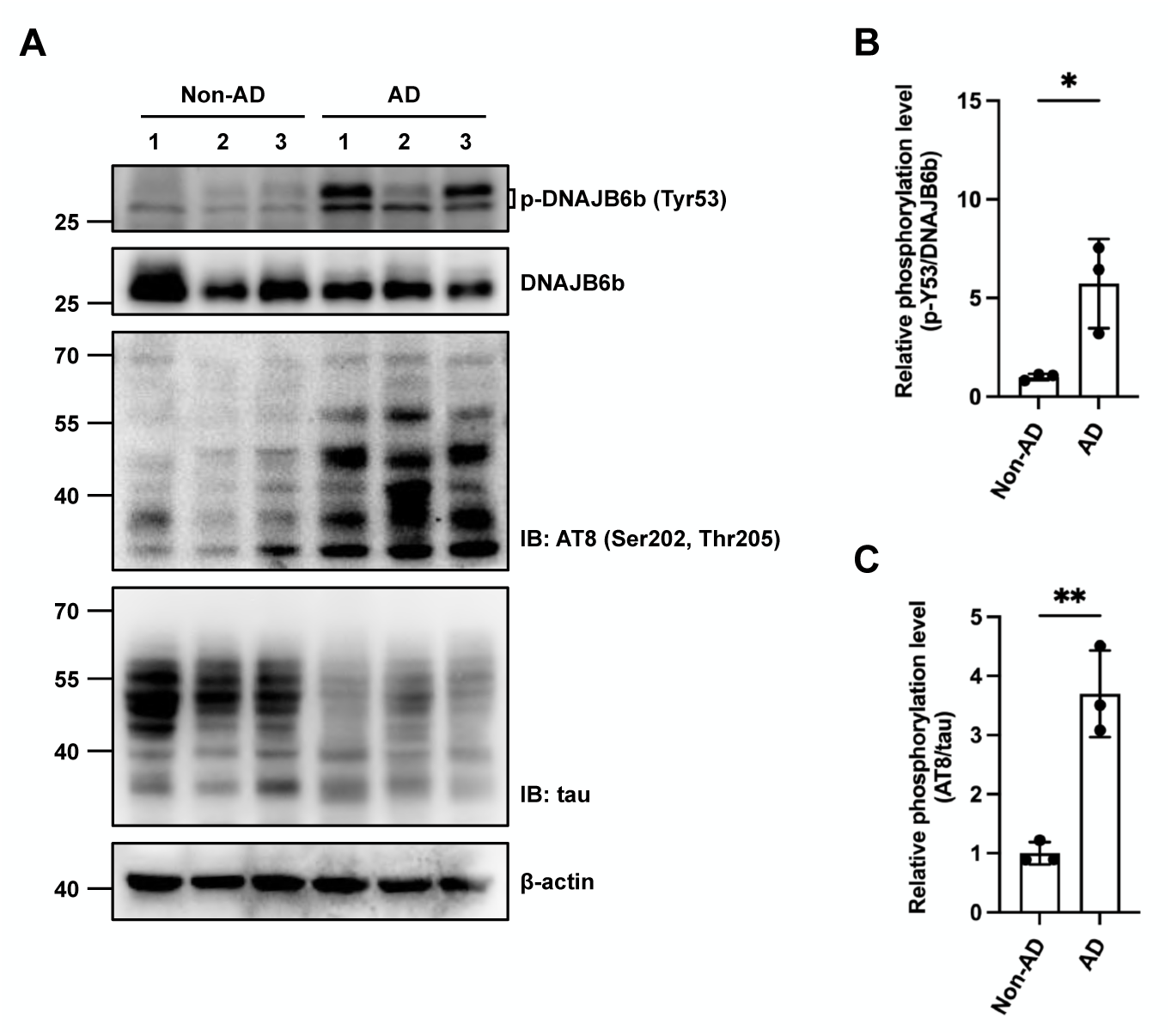
DNAJB6b Y53 phosphorylation is increased in AD brain tissues. **(A)** Commercial human whole brain lysates were acquired, and the levels of each protein were analyzed using western blotting with the specified antibodies. Phosphorylation of tau at Ser202 and Thr205 was detected using an AT8 antibody. β-actin served as a loading control. The catalog numbers for the non-AD brain lysates from 1 to 3 are #C511134, #822400674, and #C807591. The catalog numbers for the AD brain lysates from 1 to 3 are #C511136, #C511137, and #822304853. **(B)** Quantification of the ratio of phosphorylation levels of DNAJB6b Y53 compared with those of the endogenous DNAJB6b in (**A**, *n*=3). **(C)** Quantification of p-tau (Ser202 and Thr205) to tau in **(A)** (*n*=3). Data in **(B-C)** were expressed as mean ± SD, and *p*-values were calculated via unpaired two-tailed Student’s *t*-test. The individual replicate data in (**B**) and (**C**) are listed in **Table S10**. (* *p* < 0.05, ** *p* < 0.01)

Given the correlation between tau aggregation and the phosphorylation level of DNAJB6b Y53, further experiment was conducted to evaluate whether tau expression per se causes DNAJB6 to modify its activity through posttranslational modification. The Y53 phosphorylation level is evaluated in SH-SY5Y cells overexpressing EGFP-tau wild-type and P301L mutant. As shown in **Figure S2A,** the EGFP-tau P301L-overexpressed group exhibited a significantly lower Y53 phosphorylation level. On the other hand, the EGFP-tau wild-type-overexpressed group showed no differences from the control. Together, the results suggest that DNAJB6b may become hypo-phosphorylated, leading to its hyperactivation, to deal with the overwhelming aggregation-prone tau proteins overexpressed in the cells.

### SFKs regulate Y53 phosphorylation on DNAJB6b in SH-SY5Y cells

To further investigate the possible kinase responsible for DNAJB6b Y53 phosphorylation, we employed the NetPhos 3.1 server [33, 34] to predict the top three candidates, including insulin receptor (INSR) kinase, SRC, and epidermal growth factor receptor (EGFR) kinase (see **Table S2**). After cells were treated with the respective kinase inhibitors, we examined the phosphorylation levels of DNAJB6b Y53 and kinase activity compared to the solvent control. First, V5-DNAJB6b-overexpressed cells were incubated with 10 μM NVP-AEW 541, a specific inhibitor of INSR kinase (**Figure 3A-C**). The suppression of AKT serine 473 phosphorylation served as an indicator of kinase activity (**Figure 3C**) [35, 36]. The results indicated no significant difference in p-Y53 level between the wild-type group treated with and without NVP-AEW 541 (**Figure 3B**), suggesting that INSR kinase is not responsible for the phosphorylation of DNAJB6b at Y53.

**Figure 3.**
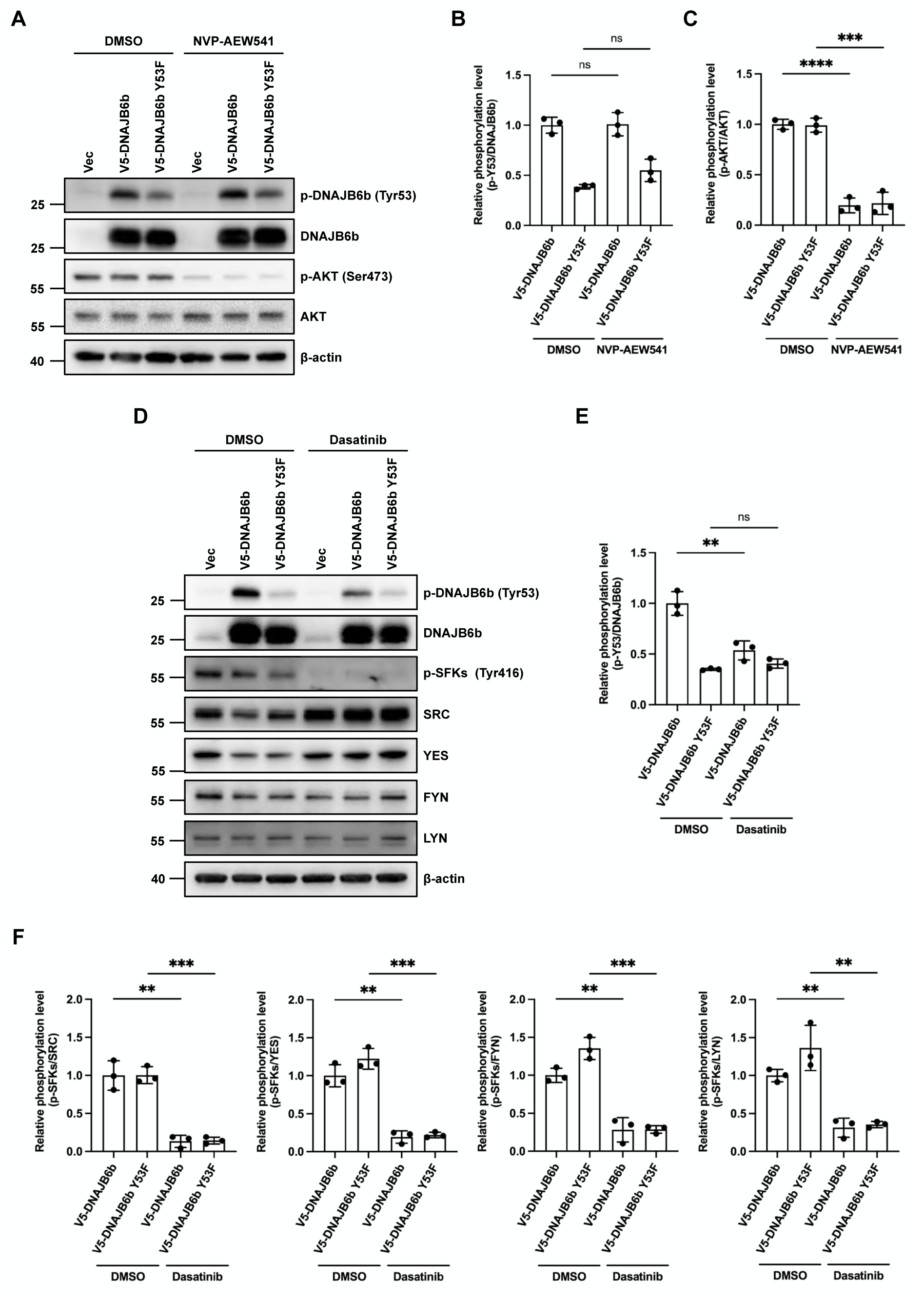
SFKs mediate the phosphorylation of DNAJB6b at Y53. **(A)** The phosphorylation of DNAJB6b at the Y53 site was evaluated in NVP-AEW541-treated SH-SY5Y cells by western blotting. Cells were over-expressed with empty vector (pcDNA/FRT/TO-V5) and indicated V5-DNAJB6b constructs for 48 hours, followed by NVP-AEW541 treatment or solvent control DMSO for 2 hours. Proteins were analyzed by immunoblotting using the indicated antibodies. **(B-C)** Quantification of the ratio of p-DNAJB6b Y53 to DNAJB6b wild-type or DNAJB6b Y53F **(B)** and the ratio of p-AKT Ser473 to AKT **(C)** in **(A)** (*n*=3). **(D)** The phosphorylation level of DNAJB6b Y53 was evaluated in dasatinib-treated cells by western blotting. SH-SY5Y cells were overexpressed with empty vector (pcDNA/FRT/TO-V5) and indicated V5-DNAJB6b constructs for 48 hours, followed by dasatinib treatment or DMSO solvent control for 2 hours. Proteins were analyzed by immunoblotting using the indicated antibodies. **(E-F)** Quantification of the ratio of p-DNAJB6b Y53 to DNAJB6b wild-type or DNAJB6b Y53F **(E)** and the ratio of p-SFKs Tyr416 to SRC, YES, FYN, or LYN **(F)** in **(D)** (*n*=3). Data in **(B-C, E-F)** were expressed as mean ± SD, and *p*-values were calculated via unpaired two-tailed Student’s *t*-test. The individual replicate data in (**B**), (**C**), (**E**), and (**F**) are listed in **Table S10**. (ns, not significant; ** *p* < 0.01, *** *p* < 0.001, **** *p* < 0.0001)

Next, we examined the second candidate, SFKs. Human SFKs comprise 11 members and can be categorized into three subfamilies based on their homology [37, 38]. The SRC subfamily (SRC, FYN, YES, and FGR) and LYN subfamily (LYN, HCK, LCK, and BLK) show similar modes of regulation in cellular viability, proliferation, differentiation, and migration [39, 40]. The SFK-associated subfamily (BRK, FRK, SRM) lacks C-terminal regulatory tyrosine and N-terminal myristoylation sites and is highly expressed in gastric and breast cancer [41, 42]. The major activation of SFKs is often led by the highly conserved autophosphorylation site, tyrosine 416 [43], which serves as an indicator of the SFKs’ activities in the following experiments. To investigate whether SFKs regulate DNAJB6b Y53 phosphorylation, SH-SY5Y cells were incubated with 500 nM dasatinib, a pan-inhibitor of SFKs [44, 45] (**Figure 3D**). Under the suppression of kinase activity, we observed a decrease in the p-Y53 level only in the V5-DNAJB6b wild-type group but not in the V5-DNAJB6b Y53F mutant (**Figure 3E**). This suggests that SFKs may regulate the phosphorylation of DNAJB6b at Y53. An additional SFK inhibitor, pyrazolopyrimidine 2 (PP2), which strongly inhibits the kinase activity of FYN, LCK, and HCK [46, 47], was also tested in SH-SY5Y cells (see **Figure S3A-C**). Interestingly, after assuring the suppression of kinase activity, the results showed no significant difference in p-Y53 level in the V5-DNAJB6b wild-type group (see **Figure S3B-C**). Together, we predicted that the upstream kinase(s) to phosphorylate DNAJB6b at Y53 may include SRC, LYN, and YES kinase, instead of FYN, LCK, and HCK kinase, in SH-SY5Y cells.

### YES kinase directly promotes the phosphorylation of DNAJB6b Y53

To better understand the distribution of individual SFKs in human tissue, particularly in the brain, the gene expression levels were analyzed. The gene expression data were sourced from the GTEx portal databank (https://www.gtexportal.org/home/, updated as of April 2, 2025) and the BrainSpan project (https://www.brainspan.org/static/home, updated as of April 2, 2025), respectively. In the GTEx portal databank, 54 non-diseased tissues from 19788 samples were examined, revealing that SRC, YES, and FYN are widely expressed across different tissues, while the others exhibited a more specific distribution (see **Figure S4A**). Additionally, five brain regions related to tau deposition [48, 49], including the medial prefrontal cortex (MFC), the superior temporal cortex (STC), the inferolateral temporal cortex (ITC), the hippocampus (HIP), and the inferior parietal cortex (IPC), were selected to further assess in seven non-diseased individuals (see **Figure S4B-C**). The findings indicated that SRC, YES, FYN, and LYN (highlighted in red) exhibit higher gene expression levels than other SFKs in these brain regions (see **Figure S4B**). Several studies have demonstrated that these SFKs may redundantly regulate neuronal survival [50], microglial migration [51], and glial phagocytic activity [52]. In **Figure S4D**, we rearranged the gene expression levels from the heatmap into a line plot. Each dot represents the z-score of gene expression in a specific brain region of the individual. The green line shows the gene expression level of FYN kinase, which remains constant across all ages. Interestingly, the gene expression levels of YES and LYN kinases follow a similar pattern, with a gradual increase before age 11 and a sudden drop at age 30. In contrast, the red line represents the gene expression level of SRC kinase, which displays a slightly opposite pattern to that of YES and LYN kinases. To conclude, SRC, YES, and LYN kinases may act as complementary players in signaling pathways at different ages.

To verify the specific SFKs responsible for DNAJB6b Y53 phosphorylation, RNAi knockdown was performed in SH-SY5Y cells by using shSRC, shYES, shFYN, and shLYN expression plasmids. Knockdown efficiencies of SFKs were examined by western blotting (**Figure 4A-L**). Compared to the knockdown of the firefly luciferase (Luc) control, only the knockdown of YES kinase (**Figure 4D**), but not SRC (**Figure 4A**), FYN (**Figure 4G**), or LYN (**Figure 4J**), led to a decrease in p-Y53 level. Knockdown of the four SFKs was also performed in other neuroblastoma, BE(2)-M17 cells (see **Figure S5A-L**). Compared to the knockdown of Luc control, a significant reduction of p-Y53 level was observed in the knockdown of YES cells (**Figure S5D-F**). These findings indicate that YES kinase plays a major role in regulating the phosphorylation of DNAJB6b Y53 in neuroblastoma cells.

**Figure 4.**
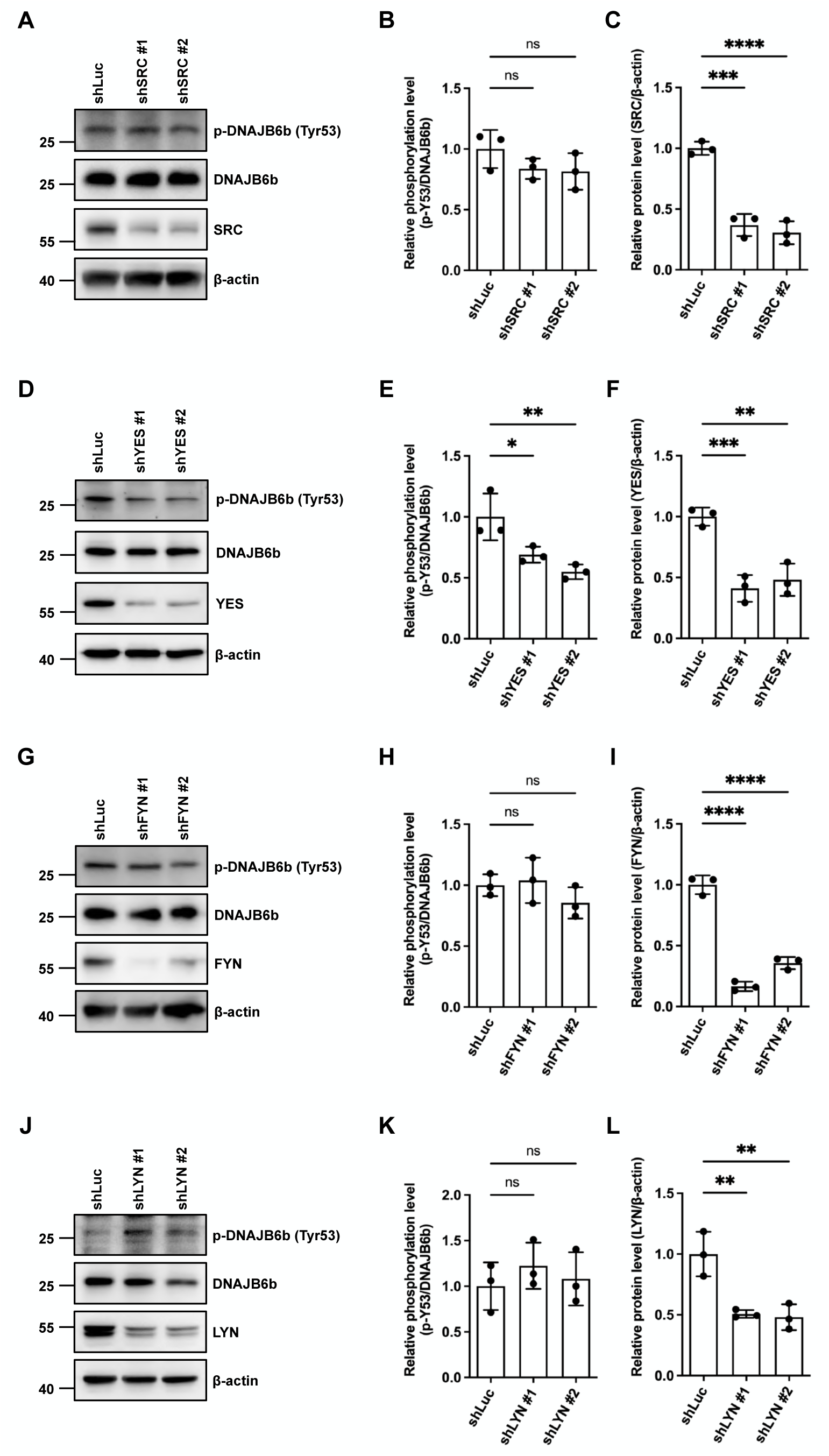
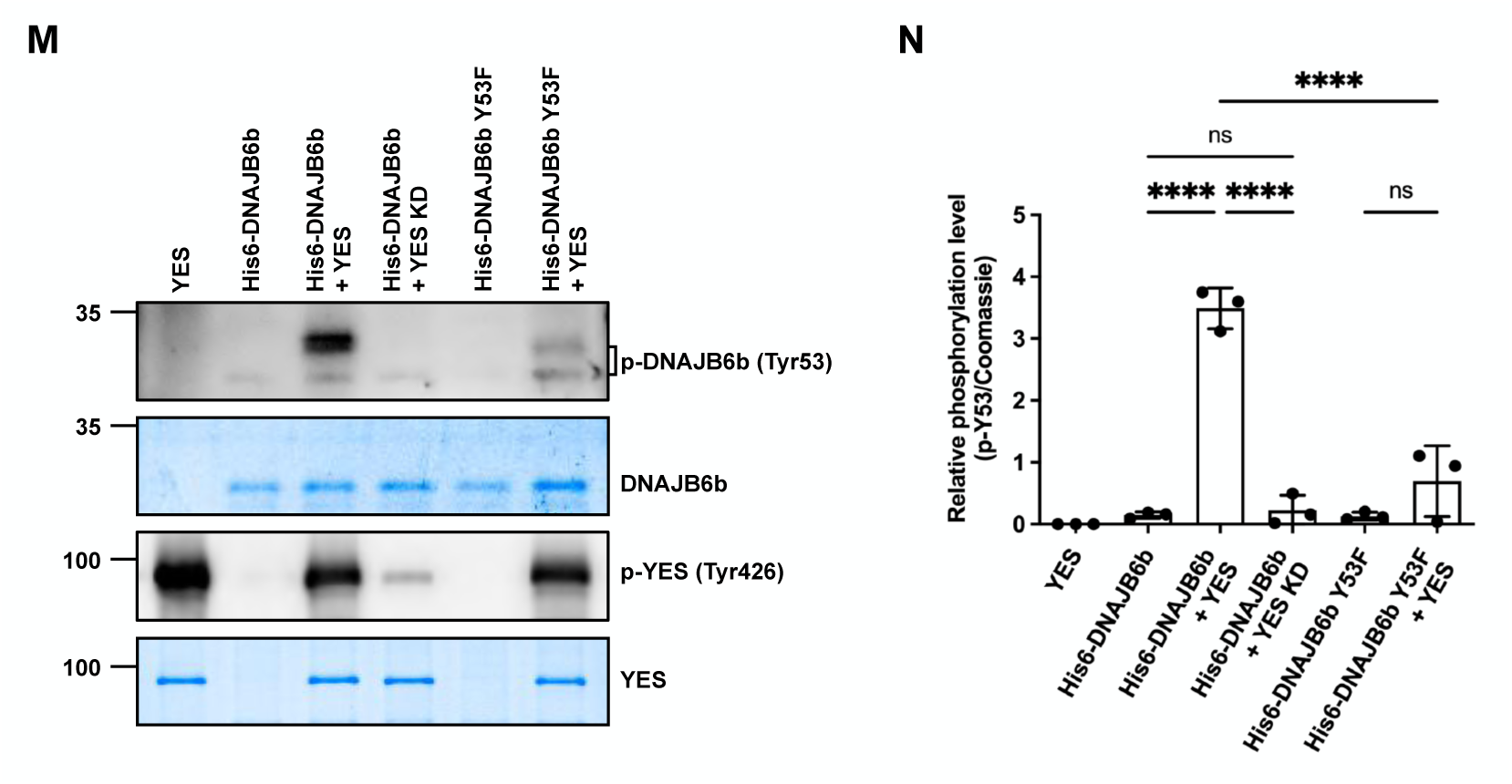
YES, but not SRC, FYN, or LYN, phosphorylates DNAJB6b Y53 in SH-SY5Y cells and in an *in vitro* kinase assay. **(A-L)** Validation of SFKs responsible for the phosphorylation of DNAJB6b Y53. Knockdown of individual SFKs in SH-SY5Y cells was performed, including SRC (**A-C**, *n*=3), FYN (**D-F**, *n*=3), LYN (**G-I**, *n*=3), and YES (**J-L**, *n*=3), and analyzed by western blotting. **(A, D, G, J)** Immunoblotting with the indicated antibodies was used to assess the phosphorylation level of DNAJB6b and the knockdown efficiency of SFKs. β-actin served as a loading control. **(B, E, H, K)** Quantification of the ratio of p-DNAJB6b Y53 to endogenous DNAJB6b in shSRC **(B)**, shFYN **(E)**, shLYN **(H)**, or shYES **(K)** groups in **(A, D, G, J)**, respectively. **(C, F, I, L)** Quantification of the ratio of SRC **(C)**, FYN **(F)**, LYN **(I)**, or YES **(L)** to β-actin in **(A, D, G, J)**. **(M)** Assessment of YES-mediated phosphorylation at Y53 on His6-DNAJB6b wild-type by *in vitro* YES kinase assay. 2 nM recombinant His6-DNAJB6b wild-type proteins were incubated with YES kinase and ATP in the suggested tyrosine kinase reaction buffer at 30°C for 15 minutes. Immunoblotting with anti-p-DNAJB6b Y53 antibody and anti-p-SFK Tyr416 antibody assessed the phosphorylation level of His6-DNAJB6b and the activity of YES kinase, respectively. Coomassie blue staining confirmed that recombinant His6-DNAJB6b and YES kinase were equally loaded. **(N)** Quantification of the ratio of p-DNAJB6b Y53 to Coomassie blue-stained His6-DNAJB6b in (**M)** (*n=3*). Data in **(B-C, E-F, H-I, K-L, N)** were expressed as mean ± SD, and *p*-values were calculated via one-way ANOVA followed by Dunnett’s test. The individual replicate data in (**B**), (**C**), (**E**), (**F**), (**H**), (**I**), (**K**), (**L**), and (**N**) are listed in **Table S10**. (ns, not significant; * *p* < 0.05, ** *p* < 0.01, *** *p* < 0.001, **** *p* < 0.0001)

To determine whether YES kinase can directly phosphorylate DNAJB6b at the Y53 site, further investigation is conducted through an *in vitro* YES kinase assay (**Figure 4M-N**). Recombinant full-length His6-DNAJB6b wild-type and His6-DNAJB6b Y53F proteins were overexpressed in *Escherichia coli*, purified, and used as substrates along with active YES kinase in the kinase assay [53] (**Figure 4M**). The presented p-Y53 band is higher than expected, which is thought to be the molecular weight shift caused by YES kinase phosphorylating other sites on DNAJB6b. The results showed a significant increase in p-Y53 level on His6-DNAJB6b wild-type incubated with active YES kinase, but not with kinase-dead (KD) YES (**Figure 4N**). Additionally, the His6-DNAJB6b Y53F mutant failed to be phosphorylated by active YES kinase. Therefore, the *in vitro* YES kinase assay demonstrates that Y53 on DNAJB6b is a direct target of YES kinase.

Next, we asked whether knockdown of YES kinase can restore the co-chaperone function of DNAJB6b and is associated with a reduction in tau aggregates. Tau aggregation levels were assessed in YES knockdown cells, followed by overexpression of EGFP-tau P301L (**Figure 5A**). The results showed that a significantly lower level of tau aggregation was observed in the YES knockdown cells, compared to the shLuc group (**Figure 5B-C**). Furthermore, with the overexpression of V5-DNAJB6b wild-type (**Figure 5D**), a significantly lower level of tau aggregation persisted in the YES knockdown groups (**Figure 5E-F**). The results suggest that, in the absence of the upstream kinase YES, DNAJB6b maximizes its co-chaperone activity and is associated with the decrease of tau aggregates accumulated in SH-SY5Y cells.

**Figure 5.**
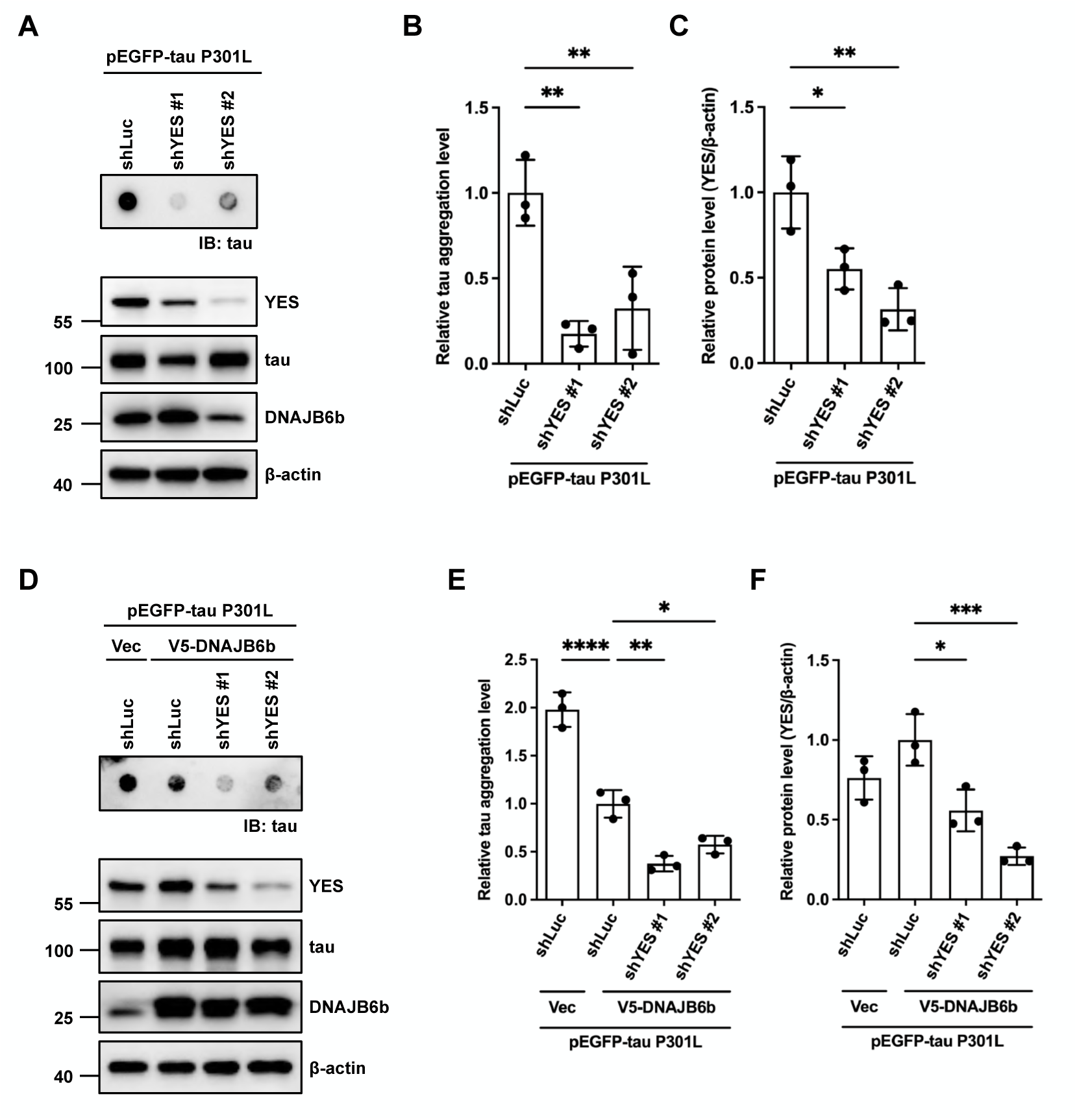
Knockdown of YES facilitates the chaperone function of DNAJB6b to prevent tau aggregation in SH-SY5Y cells. The filter trap assay for analysis and quantification of aggregated tau proteins after knockdown of YES. **(A)** After knockdown of YES, SH-SY5Y cells were co-transfected 24 hours later with EGFP-tau P301L. Retained proteins on the cellulose acetate membrane were immunoblotted using an anti-tau antibody to detect trapped tau. Immunoblotting with the indicated antibodies assessed the expression level of EGFP-tau P301L, V5-DNAJB6b, and YES. **(B)** Quantification of aggregated tau proteins. β-actin served as a loading control for normalization in **(A)** (*n*=3). **(C)** Quantification of the ratio of YES to β-actin in **(A)**. **(D)** After knockdown of YES, SH-SY5Y cells were co-transfected 24 hours later with EGFP-tau P301L and either the empty vector (pcDNA/FRT/TO-V5) or V5-DNAJB6b wild-type. Retained proteins on the cellulose acetate membrane were immunoblotted using an anti-tau antibody to detect trapped tau. Immunoblotting with the indicated antibodies assessed the expression level of EGFP-tau P301L, V5-DNAJB6b, and YES. **(E)** Quantification of aggregated tau proteins. β-actin served as a loading control for normalization in **(D)** (*n*=3). **(F)** Quantification of the ratio of YES to β-actin in **(A)**. **(B-C, E-F)** Data were expressed as mean ± SD, and *p*-values were calculated via one-way ANOVA followed by Dunnett’s test. The individual replicate data in (**B**), (**C**), (**E**), and (**F**) are listed in **Table S10**. (* *p* < 0.05, ** *p* < 0.01, **** *p* < 0.0001)

### Phosphorylation at Y53 on DNAJB6b increases the interaction with cytoplasmic Hsp70

According to previous studies, the J domain of DNAJB6b is associated with HSPA8 and HSPA1A and can enhance the binding ability with clients to resolve aggregates [21]. However, whether DNAJB6b Y53 phosphorylation can interrupt the Hsp70 machinery reaction cycle remains unclear. To evaluate whether DNAJB6b Y53 phosphorylation regulates the association with HSPA8 and HSPA1A in SH-SY5Y cells, the co-immunoprecipitation assay was further conducted (**Figure 6A**). Compared to the V5-DNAJB6b wild-type group, the V5-DNAJB6b Y53F mutant significantly reduced its interaction with Hsp70s (**Figure 6B-C**). Meanwhile, the V5-DNAJB6b Y53E mutant demonstrated a higher binding ability than both the V5-DNAJB6b wild-type and Y53F groups, indicating that phosphorylation at Y53 enhances the interaction between DNAJB6b and Hsp70s. To further examine the *in situ* protein-protein interactions of different DNAJB6b mutants with HSPA8, the PLA [54, 55] was performed (**Figure 6D-F**). Consistent with the results of the co-immunoprecipitation assay, overexpression of V5-DNAJB6b wild-type or Y53E showed a large number of PLA-positive puncta, indicating a strong interaction with HSPA8 after phosphorylation at Y53 (**Figure 6F**). Overexpression of V5-DNAJB6b Y53F displayed a relatively low PLA-positive signal, suggesting that the interaction between non-phosphorylatable DNAJB6b and HSPA8 is weak. Together, the *in vitro* cell culture along with *in situ* PLA data imply that the Y53-phosphorylated DNAJB6b establishes a strong interaction with HSPA8. The tightened DNAJB6b-Hsp70 association may result in the accumulation of tau aggregates.

**Figure 6.**
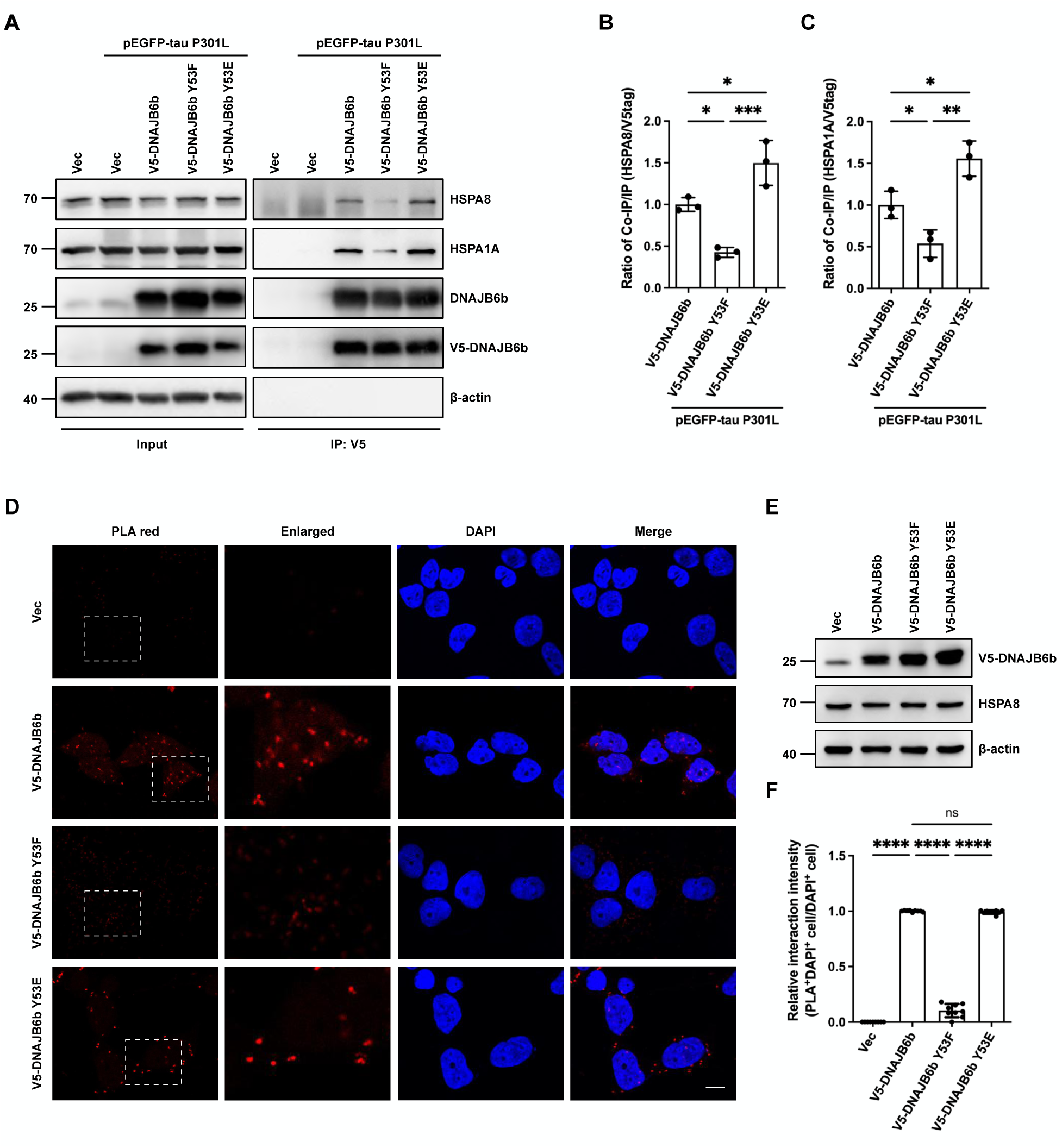
Phosphorylation of DNAJB6b Y53 strengthens the interaction between DNAJB6b and Hsp70. **(A)** Co-immunoprecipitation assay indicated the interaction between different DNAJB6b Y53 constructs and HSPA8 or HSPA1A. SH-SY5Y cells were co-expressed for 48 hours with EGFP-tau P301L and either the empty vector (pcDNA/FRT/TO-V5), V5-DNAJB6b wild-type, V5-DNAJB6b Y53F, or V5-DNAJB6b Y53E. Cell lysates were immunoprecipitated using an anti-V5 antibody to pull down V5-tagged different DNAJB6b constructs. 10% of the cell lysate from each sample is isolated to indicate the input. Precipitated and co-precipitated proteins were analyzed by immunoblotting using the indicated antibodies. **(B-C)** Quantification of the ratio of co-immunoprecipitated HSPA8 **(B)**, HSPA1A **(C)** to immunoprecipitated V5-DNAJB6b wild-type, V5-DNAJB6b Y53F, or V5-DNAJB6b Y53E in **(A)** (*n*=3). **(D)** Representative fluorescent images show the interaction between different V5-DNAJB6b constructs and HSPA8. Either the empty vector (pcDNA/FRT/TO-V5), V5-DNAJB6b wild-type, Y53F, or Y53E was transfected into SH-SY5Y cells. anti-V5 and anti-HSPA8 primary antibodies were hybridized to locate DNAJB6b and HSPA8, respectively. When different V5-DNAJB6b and HSPA8 interact, the PLA probes on secondary antibodies were ligated and amplified. The red PLA signals showing protein interaction were detected by fluorescent confocal microscopy. Nuclei were counterstained with DAPI. Scale bar: 10 μm. **(E)** Immunoblotting with anti-V5 and anti-HSPA8 antibodies indicated the expression level of different V5-DNAJB6b constructs and HSPA8, respectively. **(F)** Quantification of interaction intensity in **(D)** (*n*=3). The percentage of fluorescent (PLA^+^) SH-SY5Y cells to the total number of cells (DAPI^+^) was quantified in at least three random-field images. Data in **(B-C, F)** were expressed as mean ± SD, and *p*-values were calculated via one-way ANOVA followed by Dunnett’s test. The individual replicate data in (**B**), (**C**), and (**F**) are listed in **Table S10**. (ns, not significant; * *p* < 0.05, ** *p* < 0.01, *** *p* < 0.001, **** *p* < 0.0001)

To investigate how phosphorylation of DNAJB6b at Y53 may modulate its interaction with HSPA8, we employed AlphaFold 3 to predict the structure of the HSPA8-DNAJB6b complex in both unphosphorylated and phosphorylated states (**Figure 7**). The modeling revealed that Y53 lies within the primary dimerization interface, with the surrounding structural features predicted at high confidence (pLDDT > 90) (See **Figure S6**). In the unphosphorylated model, Y53 does not contact HSPA8 (**Figure 7A**); however, upon phosphorylation, phospho-Y53 (pY53) forms a plausible tripartite interaction with HSPA8 residues E218 and K220 (**Figure 7B**), consistent with known pTyr-Lys-Glu hydrogen-bonding geometries. This rearrangement, together with the predicted phosphorylation-induced shift of the α2-α3 helices toward HSPA8, could help stabilize the complex (**Figure 7C**) and is in line with our immunoprecipitation data showing enhanced HSPA8 binding upon Y53 phosphorylation.

**Figure 7.**
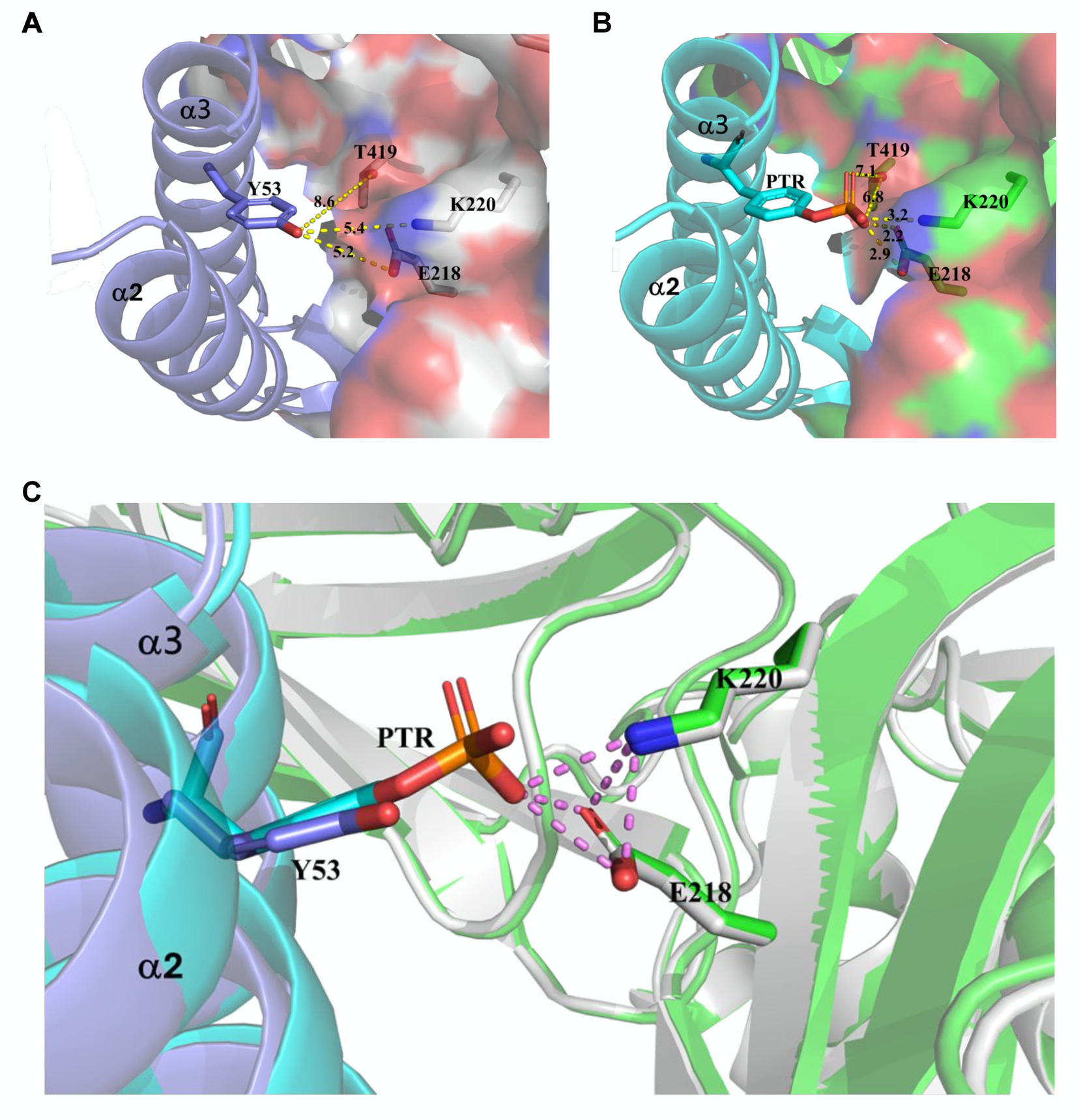
Comparison of AlphaFold 3-predicted structures of the HSPA8-DNAJB6b complex before and after DNAJB6b Y53 phosphorylation. **(A)** Predicted structure of the HSPA8-DNAJB6b dimerization interface in the unphosphorylated state. DNAJB6b is shown as purple ribbons with Y53 in stick representation. HSPA8 is displayed in surface view using grey-based CPK coloring. Residues within 12.0 Å of Y53 (E218, K220, and T419) are highlighted, with yellow dashed lines indicating their shortest distances to Y53 (measured in Å). **(B)** Predicted structure of the dimerization interface upon DNAJB6b Y53 phosphorylation. DNAJB6b is shown as cyan ribbons with phosphorylated Y53 (pY53, PTR) in stick representation. HSPA8 is rendered in surface view using green-based CPK coloring. Residues within 12.0 Å of pY53 (E218, K220, and T419) are highlighted, and yellow dashed lines indicate their shortest distances to pY53 (in Å). **(C)** Superimposed structures of the HSPA8-DNAJB6b complex before and after Y53 phosphorylation. The phosphate group of pY53 forms a hydrogen bond network (violet dashed lines) with K220 and E218 of HSPA8. Coloring follows the scheme in panels **(A)** and **(B)**.

## Discussion

In neurodegenerative diseases, loss of proteostasis is widely recognized as the primary hallmark of aging, leading to the accumulation of misfolded, hyperphosphorylated, or highly modified protein aggregates [2]. The Hsp70 chaperone system, with the assistance of JDPs, plays a crucial role in regulating the proteostasis network. In our previous findings, DNAJB6b from the DNAJB subfamily exhibits a high binding ability with HSPA8 and facilitates the untangling of tau aggregates [21]. However, it remains unclear what the role of PTM is in mediating the co-chaperone functions of DNAJB6b, and its potential association with the pathogenesis of AD.

In this study, dephosphorylation of DNAJB6b Y53 enhances its chaperone function, which is associated with a decrease in tau aggregates. Through the immunoprecipitation assay, we observed an increase *in situ* interaction between DNAJB6b Y53E and Hsp70s, especially HSPA8 and HSPA1A. The Y53 phosphorylation may influence the autoinhibitory helix V, reported to restrict access of the HPD motif to Hsp70. The predicted conformational changes near the α2-α3 region could allosterically perturb helix V positioning, thereby relieving this auto-inhibition and facilitating HSPA8 engagement. These possibilities are presented as structural hypotheses consistent with our functional observations (**Figure 6A-C**) but await future biochemical validation. A potential regulation is that YES-mediated phosphorylation of DNAJB6b Y53 may interfere with auto-inhibition on helix V, which facilitates Hsp70 engagement [56]. Therefore, the phosphorylation status of DNAJB6b may have a similar impact as mutations in DNAJB6 that cause limb-girdle muscular dystrophy. When phosphorylated, DNAJB6b’s function may remain intact; however, it might sequester Hsp70s away from other proteins that are essential for clearing aggregated tau.

Aging is a significant risk factor for AD, potentially due to cellular and molecular changes associated with advancing age [57]. One contributing mechanism may involve the increased activation of receptor tyrosine kinases (RTKs) and the SFKs pathways during aging. These pathways are critical for promoting cellular proliferation and survival, which are advantageous for aged cells and tissues [57–59]. Even though an increased Y53 phosphorylation level in the AD patient brain lysate is observed, the functional impact on DNAJB6b cannot be evaluated in this state. However, we hypothesize that the enhanced activity of RTKs and SFKs comes at a cost: the downregulation of the DNAJB6b-mediated Hsp70 folding pathway, an essential component of proteostasis. This compromise in proteostasis increases the likelihood of tau protein aggregation and Aβ plaque formation, hallmark features of AD pathology. Consequently, while aged cells benefit from improved survival mechanisms, they become more susceptible to the molecular dysfunctions underlying AD progression.

## Conclusions

Overall, this study shows that dephosphorylation at Y53 of DNAJB6b is associated with the reduction of tau aggregates. The links between Y53-phosphorylated DNAJB6b and the levels of pathological phospho-tau in AD brain support our conclusion that Y53-phosphorylated DNAJB6b may malfunction because of excessive interaction with chaperones.

## Declarations of interests

### Ethics approval and consent to participate

All animal studies were performed according to the guidelines and approved by the Institutional Animal Care and Use Committee at the Laboratory Animal Center, College of Medicine, National Taiwan University.

### Consent for publication

Not applicable.

### Availability of data and materials

All data are available upon reasonable request and directed to the corresponding authors. All materials generated in this study are available from the corresponding authors with a completed Materials Transfer Agreement.

## Acknowledgments

We thank Dr. Akihiko Takashima for the tau plasmid. We also thank the Image Core Facility of NTU for the technical support of immunofluorescence staining.

## Authors’ contributions

S.-C. Teng conceived the project. M.-C. Lin, S.-J. Lin and C.-Y. Liao prepared the experimental materials. M.-C. Lin and P.-J. Lin cultured the cells, prepared the protein samples, and performed the biochemical assays and BiFC assay. T.-S. Lin and N.-L. Chan conducted the protein docking studies. H.-Y. Chen performed the PLA. All authors contributed to the data analysis. M.-C. Lin, C.-H. Lin, and S.-C. Teng designed the experiments and wrote the manuscript with input from all co-authors.

## Supplemental information

Supplementary PDF. Include Figures S1–S7 and Tables S1–S9.

Tables S10. The individual data values of the replicates, related to Figures 1-6, S1–S3, S5.

## Methods

### Cell lines and culture conditions

SH-SY5Y and BE(2)-M17 neuroblastoma cells were obtained from the American Type Culture Collection (ATCC) and cultured in Dulbecco’s Modified Eagle Medium/Nutrient Mixture F-12 (DMEM/F12) medium (Cytiva, Logan, UT, USA) supplied with 10% fetal bovine serum (VWR, Radnor, PA, USA) and 1x penicillin/streptomycin/amphotericin B (Capricorn Scientific, Germany). Cells were plated on poly-L-lysine-coated 10 cm cell culture dishes or 6-well culture plates and grown in a humidified incubator at 37°C with 5% CO_2_.

### Plasmids

Primers used for plasmid generation and the specific oligonucleotide sequences of shRNA are listed in Document S1: **Table S3** and **Table S4**. Construction of pEGFP-C1-tau P301L (2N4R) and pcDNA5/FRT/TO-V5 control was performed as described [21]. The VN-tau (P301L), tau (P301L)-VC, and pcDNA5/FRT/TO-V5-DNAJB6b were purchased from Addgene (Watertown, MA, USA). pcDNA5/FRT/TO-V5-DNAJB6b Y53F and pcDNA5/FRT/TO-V5-DNAJB6b Y53E plasmids were generated by site-directed mutagenesis using HiFi polymerase (Kapa Biosystems, Wilmington, MA, USA) with DNAJB6-Y53F-BstBI-For and DNAJB6-Y53E-NdeI^del^-For, respectively. All plasmids were sequenced before use and are listed in Document S1: **Table S5**.

### Transient transfections, shRNA transfections, and sample preparation

Either transient transfections or shRNA transfections were performed using Lipofectamine™ LTX reagent with PLUS™ reagent (Thermo Fisher Scientific, Waltham, MA, USA) according to the manufacturer’s instructions. Cells were grown to 60-70% confluence on poly-D-lysine-coated 6-well plates and transfected for 48-72 hours. For shRNA transfection, cells were additionally selected with 2-4 μg/ml of puromycin (Sigma-Aldrich, Burlington, MA, USA) for at least 2 days. Cell pellets were collected and lysed on ice for 10 minutes in RIPA lysis buffer (Merck Millipore, Darmstadt, Germany), followed by centrifugation at 13,000 rpm for 15 minutes at 4°C. The supernatants were collected, and the protein concentration was determined by the Bio-Rad Protein Assay (Bio-Rad Laboratories, Hercules, CA, USA). 5x SDS sample buffer (250 mM Tris-HCl, pH 6.8, 10% SDS, 0.25% bromophenol blue, 50% glycerol, 10 mM DTT, 0.5 M 2-mercaptoethanol) was then added to the supernatants, and the samples were heated at 110 °C for 15 minutes. Samples were stored at -80°C for further analysis.

### Filter trap assay

The filter trap assay was performed as previously described [21, 60]. For the DNAJB6 wild-type and tau P301L groups, cells were co-transfected with 1.5 µg and 0.75 µg of plasmids, respectively. For the DNAJB6b mutant and tau P301L groups, 1.5 µg and 0.5 µg of plasmids were used. Cell pellets were collected and lysed with 1 % Triton X-100 in PBS solution supplemented with 1 mM PMSF, 1x cOmplete™ EDTA-free Protease Inhibitor Cocktail (Roche, Switzerland), and 1x PhosSTOP Phosphatase Inhibitors (Roche). After a total of 20 seconds of sonication, the protein concentration was determined using Bio-Rad Protein Assay (Bio-Rad Laboratories) and diluted to a final concentration of 1 μg/μL with lysis buffer containing 1% SDS. 60-90 μL solution per sample was then passed through 0.2 μm cellulose acetate membranes (Sterlitech, USA) using a 96-well dot blot apparatus (Bio-Rad Laboratories). Before filtration, the membranes were prewashed with 0.1% SDS in PBS solution. Proteins trapped by the membranes were detected by immunostaining following the protocols described in immunoblotting.

### Immunoblotting

A total of 15-30 μg protein was resolved by SDS-polyacrylamide gel electrophoresis (SDS-PAGE) and transferred to a polyvinylidene difluoride membrane (PVDF) (Merck Millipore). The membrane was blocked in 5% nonfat milk or 5% BSA at room temperature for 1 h, followed by incubation with primary antibodies at 4°C overnight. Primary antibodies (Document S1: **Table S6**) were used to detect tau (1:1000, GeneTex, Irvine, CA, USA), GFP (1:2000, GeneTex), Phospho-DNAJB6b (Tyr53) (1:250, this study), DNAJB6 (1:2000, Abcam, Cambridge, UK), V5 (1:1000, Abcam), AT8 (1:500, Thermo Fisher Scientific), Phospho-AKT (Ser473) (1:1000, Cell Signaling Technology), AKT (1:1000, Cell Signaling Technology), Phospho-Src Family (Tyr416) (1:500, Cell Signaling Technology), SRC (1:1000, Cell Signaling Technology), FYN (1:1000, GeneTex), Phospho-LYN (1:1000, Abcam), LYN (1:1000, GeneTex), YES (1:500, Cell Signaling Technology), HSPA8 (1:1000, Novus Biologicals, Centennial, CO, USA), HSPA1A (1:1000, Santa Cruz Biotechnology, Dallas, TX, USA), and β-actin (1:2000, Proteintech, Rosemont, IL, USA). Next, the membrane was washed three times with 1x Tris-buffered saline with Tween-20 (20 mM Tris-HCl, pH 7.6, 150 mM NaCl, 0.1% Tween-20), followed by incubation with the horseradish peroxidase-conjugated goat anti-rabbit or goat anti-mouse IgGs (1:5000, Jackson ImmunoResearch, West Grove, PA, USA) at room temperature for at least 2 hours. Luminata™ Crescendo Western HRP Substrate (Merck Millipore) was used for visualization by ChemiDoc™ Imaging System (Bio-Rad Laboratories). The images were quantified by Image Lab software (Bio-Rad Laboratories).

### Bimolecular fluorescence complementation (BiFC) assay

The BiFC assay was performed according to previous studies [21, 31]. SH-SY5Y cells were co-transfected with different DNAJB6b constructs, along with plasmids that encode full-length tau P301L protein fused to the N-terminal part of Venus protein, VN-tau P301L (Addgene), and tau P301L protein fused to the C-terminal part of Venus protein, tau P301L-VC (Addgene). After 48 hours of transfection, cells were immunostained with V5-DNAJB6 (anti-V5 antibody, 1:100, Thermo Fisher Scientific) and counterstained with 4’, 6-diamidino-2-phenylindole (DAPI) (1:1000, Thermo Fisher Scientific). The transfection efficiency of the plasmids was verified by immunoblotting with an anti-DNAJB6 antibody (Abcam) and an anti-tau antibody (GeneTex), respectively. To quantify the relative Venus-tau aggregates developed in cells, three random-field images were taken at 10x magnification on a Zeiss ApoTome 2 fluorescence microscope. Three independent experiments were performed. Final representative images were taken at 40x magnification. Images were viewed using Zen Blue 2.6 (Zeiss) and analyzed using ImageJ (NIH).

### Preparation of phosphor-DNAJB6b (Tyr53) polyclonal antibodies

Synthesized peptide with a length of 12 a.a. and an additional cysteine on the C-terminal for Keyhole Limpet Hemocyanin (KLH) conjugation: A-E-A-(pY)-E-V-L-S-D-A-K-K-C (MW: 1506.57) is ordered from Synbio technologies (NJ, USA). KLH conjugation was performed utilizing the ImjectTM Maleimide-Activated mcKLH Spin Kit following the protocols prescribed by the manufacturer (Thermo Fisher Scientific). 50 μl of the peptide-KLH conjugates was administered intraperitoneally into a naïve BALB/c mouse. Following a period of three weeks, the previously primed mouse received two booster injections with the peptide-KLH. The affinity and specificity of phosphor-DNAJB6b Tyr53 polyclonal antibodies were identified through Western blotting.

### Human brain lysate samples

Non-AD control (NB820-59177, GTX27918) and AD (NB820-59363, GTX26622) post-mortem whole brain lysates were purchased from Novus Biologicals and Genetex. For biological replicates, products from different individuals were used. Product #NB820-59177 includes the non-AD brain lysate of a 26-year-old Asian male (lot #C511134) and an 82-year-old Caucasian male (lot #C807591), both of whom had no clinical evidence of neurological disorders. Product #GTX28771 is obtained from an 82-year-old Caucasian male (lot #822400674), who has known pathologies of aortic stenosis. Products #NB820-59363 and #GTX26622 contain brain lysates from individuals with confirmed Alzheimer’s disease (AD) pathology, including a 93-year-old Hispanic female (Lot #C511136), an 87-year-old Caucasian male (Lot #C511137), and a 93-year-old female (Lot #822304853). All the details of the individuals are listed in Document S1: **Table S7**.

### Preparation of stock solutions and treatment conditions

NVP-AEW541, dasatinib, and pyrazolopyrimidine (PP)2 were purchased from MedChemExpress (Monmouth Junction, NJ, USA) for inhibitor treatments. For standardizing the dose of NVP-AEW541, SH-SY5Y cells were treated for 2 hours with NVP-AEW541 or solvent control DMSO (Sigma-Aldrich) at doses of 1 μM, 10 μM, or 50 μM [35]. The lowest dosage affecting insulin receptor kinase activity levels was assessed, and the final concentration of 10 μM NVP-AEW541 was applied. Insulin receptor kinase activity was determined through the phosphorylation of AKT on serine 473.

For inhibiting the kinase activity of SFKs, cells were treated for 1 hour with 500 nM or 750 nM of the pan-inhibitor dasatinib [44] and assessed for the lowest dosage affecting SFK activity levels. For the selective inhibitor PP2, cells were treated for 6 hours with a final concentration of 5 μM, 10 μM, or 20 μM, based on previous studies [46, 61]. The final concentration of 500 nM dasatinib and 10 μM PP2 was employed, and SFKs activity was determined through the phosphorylation of SFKs on tyrosine 416.

### Expression analysis

The tissue gene expression data were obtained from the Genotype Tissue Expression (GTEx) portal database (https://www.gtexportal.org/home/) on April 2^nd^, 2025. The GTEx Multi-Gene Query platform was utilized to investigate the gene expression levels of 11 SFKs, including SRC, YES1, FYN, FGR, LYN, HCK, LCK, BLK, PTK6 (BRK), FRK, and SRMS (SRM), in 54 non-diseased tissues from 19,788 samples. The TPM (Transcripts per million) metric was used to quantify gene expression levels.

The gene expression data of 11 SFKs and DNAJB6 from five brain regions, the medial prefrontal cortex, the superior temporal cortex, the inferolateral temporal cortex, the hippocampus, and the inferior parietal cortex, were obtained from the BrainSpan project database (https://www.brainspan.org/static/home) on April 2^nd^, 2025. The log_2_ RPKM (reads per kilobase per million) values in seven individuals at different ages (12 pcw, 24 pcw, 1, 11, 21, 30, and 37 years old) were downloaded and converted into z-scores for visualization in a heatmap by the following equation:

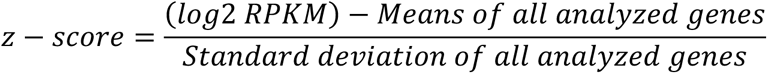

The original search can be reproduced on the BrainSpan website at: https://www.brainspan.org/rnaseq/searches?search_type=user_selections.

### Recombinant His6-DNAJB6b purification and *in vitro* YES kinase assay

The recombinant full-length His6-DNAJB6b WT and Y53F were constructed by the T4 ligation method. pcDNA5/FRT/TO-V5-DNAJB6b WT and Y53F were digested with BamHI-HF and NotI-HF as “insert” and ligated to the restriction enzyme-digested pET-28a vector with T4 ligase. Then, the recombinant His6-DNAJB6b WT and Y53F was produced in BL21(tRNA) bacteria, followed by IPTG induction, and purified by affinity to histidine-tagged protein purification resin (Cytiva). Protein concentration was determined by Bio-Rad Protein Assay (Bio-Rad Laboratories). The purified His6-DNAJB6b was submitted to buffer exchange by using an Amicon Ultra-0.5 Centrifugal Filter Unit (3 kDa, Merck Millipore) and stored in 1x tyrosine kinase reaction buffer (20 mM HEPES pH 7.4, 1 mM MnC1_2_, 1 mM DTT, 100 µM Na_3_VO_4_) with 20 % glycerol at -80°C.

For *in vitro* YES kinase assays, 2 nM His6-DNAJB6b or His6-DNAJB6b Y53F was incubated with 10 ng active GST-YES kinase (Thermo Fisher Scientific) and 200 µM ATP in tyrosine kinase reaction buffer for 15 minutes at 30°C. The reactions were quenched by the addition of 7.5 µL 5x sample buffer and boiled at 110°C for 10 minutes. Samples were analyzed by western blotting, and the homogeneity of the recombinant His6-DNAJB6b was checked by Coomassie blue staining.

### Co-immunoprecipitation assay

SH-SHY5Y cells were harvested and lysed for 30 minutes in immunoprecipitation (IP) buffer (50 mM Tris–HCl pH 7.5, 150 mM NaCl, 0.5% Triton X-100, 1 mM EDTA), supplemented with 1 mM PMSF, 1x Complete EDTA-free Protease Inhibitor Cocktail (Roche), and 1x PhosSTOP Phosphatase Inhibitors (Roche). 2 μg Anti-V5 antibody (Abcam) was mixed with a final concentration of 2 μg/ml lysate and incubated tumbled end-over-end for 1 hour at 4°C. Lysates were then incubated with Protein G Mag Sepharose Xtra magnetic beads (Cytiva) for 90 minutes at 4°C. After extensive washing with wash buffer (50 mM Tris-HCl, pH 7.5, 150 mM NaCl, 0.5% Triton X-100, 1 mM EDTA) three times, the bound proteins were eluted by the addition of 30 μL 2x SDS sample buffer. Precipitates were boiled for 10 minutes at 110°C before analysis by SDS-PAGE.

### Proximity Ligation Assay (PLA)

To determine the *in situ* interaction between different DNAJB6b constructs and HSPA8, PLA was performed using the Duolink® *In Situ* Red Starter Kit Mouse/Rabbit (Sigma-Aldrich), according to the manufacturer’s protocol. SH-SY5Y cells were seeded at 1 × 10^4^ cells/ml onto circular glass coverslips (Marienfeld Laboratory Glassware, Germany) in 12-well plates and were transfected with different DNAJB6b constructs. After transfection for 48 hours, cells were fixed with 4% paraformaldehyde in PBS solution for 15 min at room temperature, followed by permeabilization with 0.1% Triton X-100 in PBS solution for 5 min. The coverslips were then blocked with 1x Duolink blocking solution for 30 min and incubated with anti-V5 (1:50, Thermo Fisher Scientific) and anti-HSPA8 (1:250, Novus) antibodies in Duolink antibody diluent at 4°C overnight. After incubation, coverslips were washed twice with Duolink Wash Buffer A and incubated with both PLA PLUS and PLA MINUS probes in a 1:5 dilution for 1 hour at 37°C. Ligation (1:40 in ligase solution) was carried out for 30 minutes, followed by amplification (1:80 in amplification solution) for 100 minutes using Detection Reagents Red. Afterward, coverslips were washed with Duolink Wash Buffer B, and the nuclei were stained with DAPI (1:1000, Thermo Fisher Scientifics). Coverslips were later mounted with 5 mL of Duolink *in situ* mounting medium and imaged using a Zeiss LSM880 confocal microscope.

To quantify the relative interaction intensity (PLA^+^DAPI^+^/ DAPI^+^ cells), 3 random-field images were acquired at 40x magnification for each condition using the Zeiss LSM880 fluorescence microscope. Three independent experiments were performed. Final representative images were acquired at 63x magnification. Images were viewed using Zen Blue 2.6 (Zeiss) and analyzed using ImageJ (NIH).

### Protein docking studies

The full-length amino-acid sequences of human HSPA8 (UniProt E9PS65) and DNAJB6b (UniProt O75190) were submitted to the AlphaFold 3 server (https://alphafoldserver.com/welcome) for multimeric complex prediction. Two models of DNAJB6b were generated: one with unmodified Y53 and another with Y53 phosphorylated. The resulting unphosphorylated and phosphorylated complexes were imported into the PyMOL Molecular Graphics System (Version 3.1.3.1; Schrödinger, LLC) for structural analysis and superposition. Cα atoms corresponding to residues 1–500 of HSPA8 were aligned using the *super* command, yielding an overall root-mean-square deviation (RMSD) of 0.25 Å.

### Statistical analysis

The protein levels in western blotting and filter trap assays were quantified using Image Lab software (Bio-Rad Laboratories). All data are presented as mean values ± standard deviation (SD), calculated using Numbers version 14.3 (Apple, CA, USA) and presented using GraphPad Prism 9 version 9.5.1 (GraphPad Software, MA, USA). All experiments were performed in a minimum of three biological replicates. The statistical test performed for each experiment was: an unpaired two-tailed Student’s t-test or one-way ANOVA, followed by Dunnett’s test for multiple comparisons. *P*-values < 0.05 were considered statistically significant and were indicated in the figures. (ns, not significant; * *p* < 0.05; ** *p* < 0.01; *** *p* < 0.001; **** *p* < 0.0001).

## List of Abbreviations

AD: Alzheimer’s disease
BiFC: Bimolecular fluorescence complementation
SFKs: Src family kinases
MAP: Microtubule-associated protein
NFTs: Neurofibrillary tangles
HSP: Heat shock protein
JDPs: J-domain proteins
NEF: Nucleotide exchange factors
TTR: Transthyretin
APP: Amyloid precursor protein
PTM: Posttranslational modification
ATCC: American Type Culture Collection
DMEM: Dulbecco’s modified Eagle’s medium
SDS-PAGE: SDS-polyacrylamide gel electrophoresis
PVDF: Polyvinylidene difluoride membrane
DAPI: 4’, 6-Diamidino-2-phenylindole
KLH: Keyhole Limpet Hemocyanin
PP2: pyrazolopyrimidine 2
EGFR: epidermal growth factor receptor
TPM: Transcripts per million
RPKM: Reads per kilobase per million
shLuc: Knockdown of firefly luciferase
IP: Immunoprecipitation
PLA: Proximity ligation assay
SD: Standard deviation
RTK: Receptor tyrosine kinase

