## Supplemental PDF for "Dephosphorylation of YES kinase-mediated co-chaperone DNAJB6b phosphorylation attenuates tau protein aggregation"

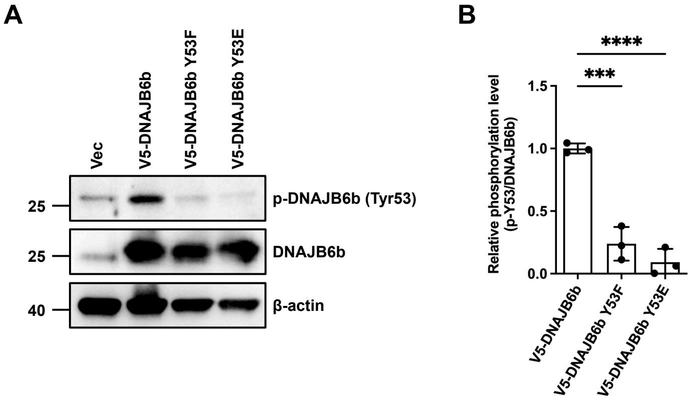


**Figure S1. Validation of phosphor-specific antibody of DNAJB6b Tyr53.**

**(A)** Validation of anti-p-DNAJB6b Y53 antibody using western blotting. SH-SY5Y cells were transfected with either the empty vector (pcDNA/FRT/TO-V5), V5-DNAJB6b wild-type, Y53F, or Y53E. The expression of different V5-DNAJB6b constructs was assessed by immunoblotting with DNAJB6 antibodies. β-actin served as a loading control. **(B)** Quantification of the ratio of p-DNAJB6b Y53 to DNAJB6b wild-type, Y53F, or Y53E in (**A)** ($n$=3). Data were represented as mean ± SD, and $p$-values were calculated via one-way ANOVA followed by Dunnett´s test. The individual replicate data are listed in **Table S10**. (*** $p < 0.001$, **** $p < 0.0001$)

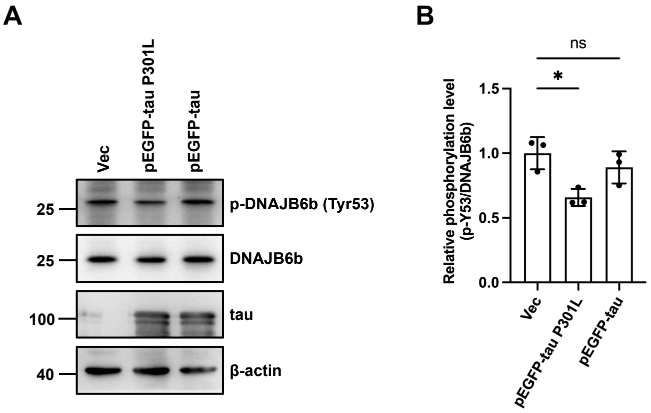


**Figure S2. Overexpression of EGFP-tau P301L downregulates the phosphorylation of DNAJB6b Y53 in SH-SY5Y cells.**

**(A)** The phosphorylation of DNAJB6b at the Y53 site was evaluated in SH-SY5Y cells by western blotting. SH-SY5Y cells were transfected with either the empty vector (pEGFP-C1), EGFP-tau P301L, or wild type. The expression of different EGFP-tau constructs was assessed by immunoblotting with tau antibodies. β-actin served as a loading control. **(B)** Quantification of the ratio of p-DNAJB6b Y53 to DNAJB6b wild-type in (**A**) ($n$=3). Data were represented as mean ± SD, and $p$-values were calculated via one-way ANOVA followed by Dunnett´s test. The individual replicate data are listed in **Table S10**. (ns, not significant; * $p < 0.05$)

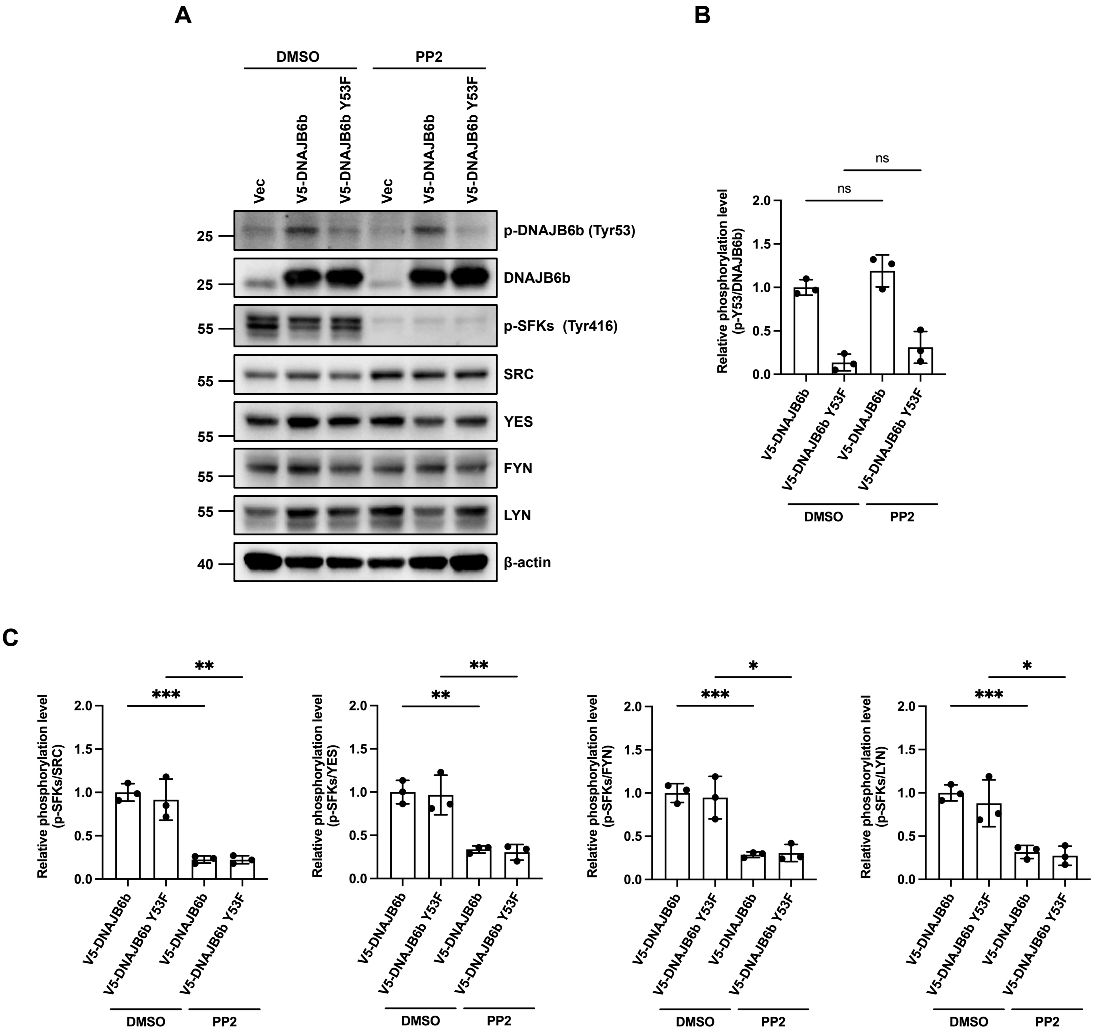


**Figure S3. Src family kinases, other than FYN, LCK, and HCK, mediate the phosphorylation of DNAJB6b at Y53. (Related to Figure 3)**

**(A)** The phosphorylation level of DNAJB6b Y53 was evaluated in PP2-treated cells by western blotting. SH-SY5Y cells were overexpressed with empty vector (pcDNA/FRT/TO-V5) and indicated V5-DNAJB6b constructs for 48 hours, followed by 10 μM PP2 treatment or DMSO solvent control for 6 hours. Proteins were analyzed by immunoblotting using the indicated antibodies. **(B-C)** Quantification of the ratio of p-DNAJB6b Y53 to DNAJB6b wild-type or DNAJB6b Y53F **(B)** and the ratio of p-SFKs Tyr416 to SRC, YES, FYN, or LYN **(C)** in (**A**) (*n*=3). Data in (**B-C**) were expressed as mean ± SD, and *p*-values were calculated via unpaired two-tailed Student's t-test. The individual data values of the replicates in (**B**), (**C**) are listed in **Table S10**. (ns, not significant; * $p < 0.05$, ** $p < 0.01$, *** $p < 0.001$)

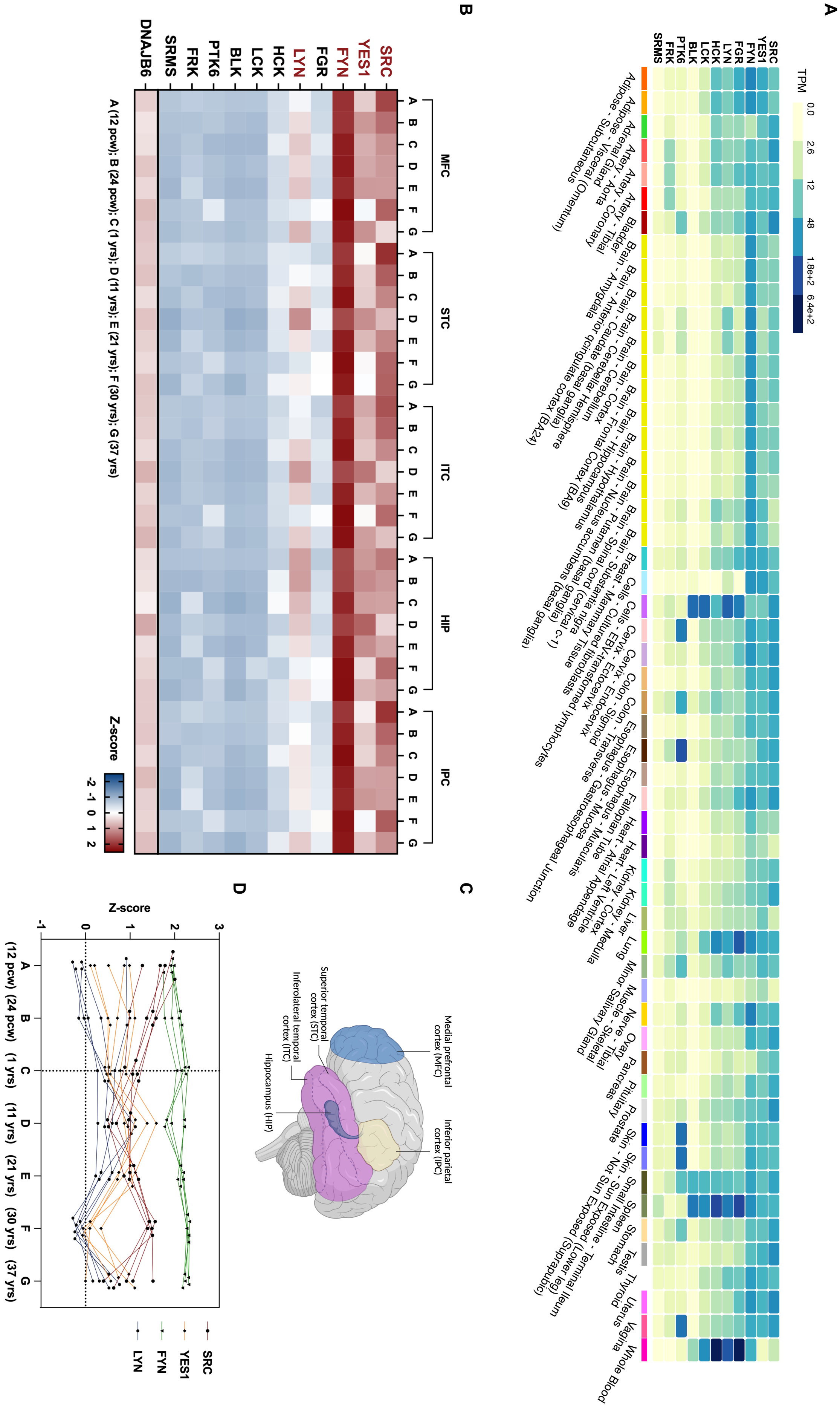

A
B
C
D
TPM
0.0
2.6
12
48
1.8e+2
6.4e+2
SRC
YES1
FYN
FGR
LYN
HCK
LCK
BLK
PTK6
FRK
SRMS
DNAJB6
MFC
STC
ITC
HIP
IPC
Z-score
A (12 pcw); B (24 pcw); C (1 yrs); D (11 yrs); E (21 yrs); F (30 yrs); G (37 yrs)
Medial prefrontal cortex (MFC)
Superior temporal cortex (STC)
Inferolateral temporal cortex (ITC)
Hippocampus (HIP)
Inferior parietal cortex (IPC)

**Figure S4. Gene expression profiles of individual SFKs in human tissues and various brain regions**

The mRNA expression of 11 Src family kinases, including SRC, YES1, FYN, FGR, LYN, HCK, LCK, BLK, PTK6 (BRK), FRK, and SRMS (SRM), in human tissues. **(A)** The heatmap presents the mRNA expression of individual SFKs in 54 non-diseased tissues, derived from 19,788 samples. Data was downloaded from the GTEx Multi-Gene Query platform from the GTEx portal databank (https://www.gtexportal.org/home/, updated as of April 2, 2025). The TPM metric was used to quantify gene expression levels. **(B)** The heatmap presents the mRNA expression of individual SFKs in five brain regions from **(C)** in seven non-diseased individuals. Data were downloaded from the BrainSpan project database (https://www.brainspan.org/static/home) on April 2$^{nd}$, 2025. The $\log_2$ RPKM values in seven individuals at different ages (12 pcw, 24 pcw, 1, 11, 21, 30, and 37 years old) were downloaded and converted into z-scores for visualization. The graph was plotted with GraphPad Prism 9 software. **(C)** Five brain regions related to tau deposition are highlighted in various colors. The medial prefrontal cortex (MFC) is marked as blue; the superior temporal cortex (STC) as the upper side of the purple region; the inferolateral temporal cortex (ITC) as the lower side of the purple region; the hippocampus (HIP) as dark blue, and the inferior parietal cortex (IPC) as light yellow. The illustration was created with BioRender.com. **(D)** The line plots present the mRNA expression pattern of four individual SFKs in five brain regions from **(C)** in seven non-diseased individuals at different ages (12 pcw, 24 pcw, 1, 11, 21, 30, and 37 years old). Data were downloaded and processed as described in **(B)**. The gene expression pattern of SRC, YES1, FYN, and LYN is marked in red, orange, green, and blue, respectively. Each dot represents the z-score of gene expression in a specific brain region of the individual. The graph was plotted with GraphPad Prism 9 software. The individual data values are listed in **Table S10**. (pcw: post conception weeks)

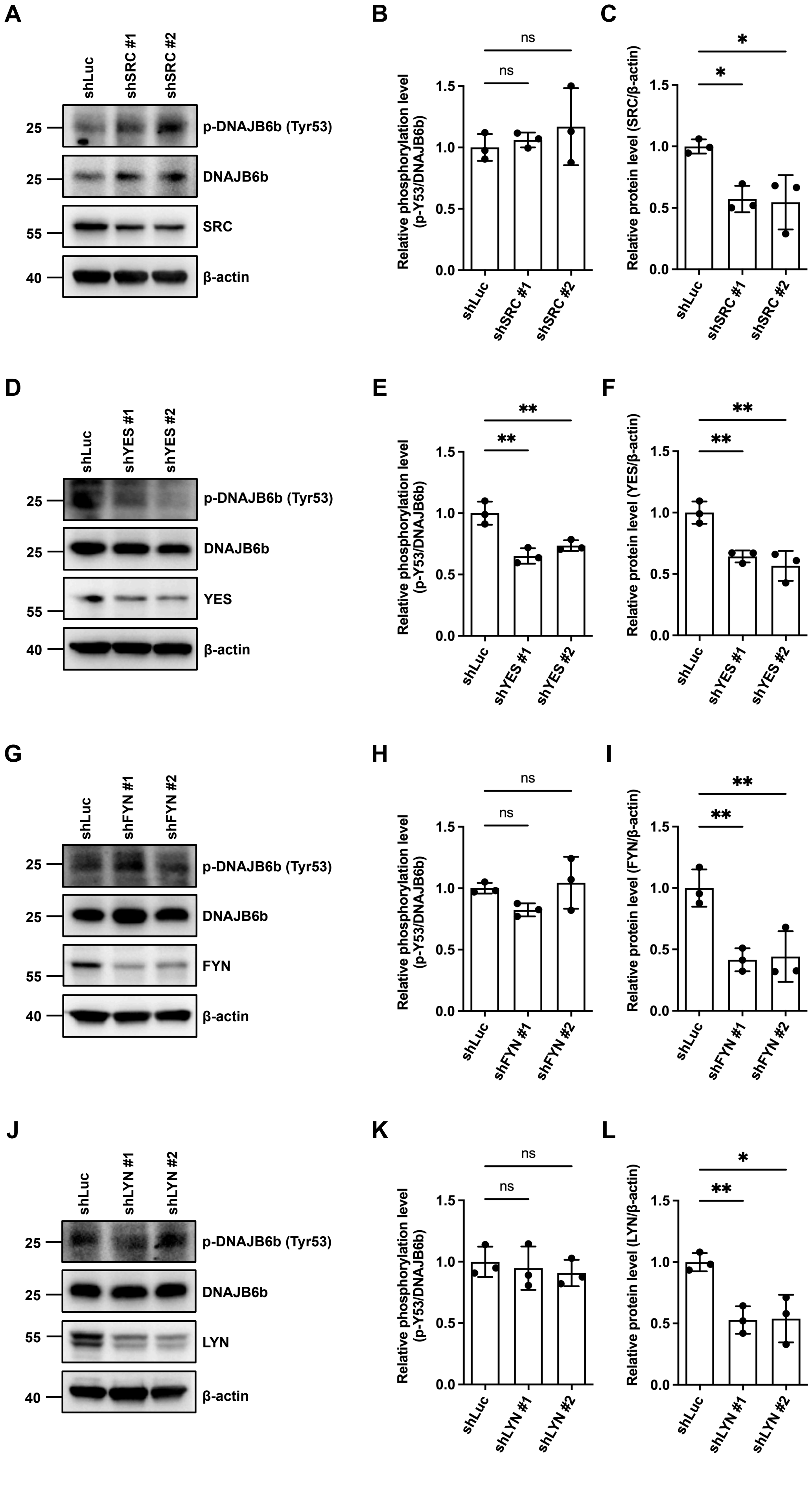
A
shLuc
shSRC #1
shSRC #2
p-DNAJB6b (Tyr53)
DNAJB6b
SRC
β-actin
B
Relative phosphorylation level (p-Y53/DNAJB6b)
C
Relative protein level (SRC/β-actin)
D
shYES #1
shYES #2
YES
E
F
Relative protein level (YES/β-actin)
G
shFYN #1
shFYN #2
FYN
H
I
Relative protein level (FYN/β-actin)
J
shLYN #1
shLYN #2
LYN
K
L
Relative protein level (LYN/β-actin)
ns
*
**

**Figure S5. YES is responsible for DNAJB6b Y53 phosphorylation in BE(2)-M17 cells, related to Figure 4**

**(A-L)** Validation of SFKs responsible for the phosphorylation of DNAJB6b Y53. Knockdown of individual SFKs in BE(2)-M17 cells was performed, including SRC (**A-C**) (*n*=3), FYN (**D-F)** (*n*=3), LYN (**G-I)** (*n*=3), and YES (**J-L)** (*n*=3), and analyzed by western blotting. **(A, D, G, J) I**mmunoblotting with the indicated antibodies was used to assess the phosphorylation level of DNAJB6b and the knockdown efficiency of SFKs. β-actin served as a loading control. **(B, E, H, K)** Quantification of the ratio of p-DNAJB6b Y53 to endogenous DNAJB6b in shSRC **(B)**, shFYN **(E)**, shLYN **(H)**, or shYES **(K)** groups in **(A, D, G, J)**, respectively. **(C, F, I, L)** Quantification of the ratio of SRC **(C)**, FYN **(F)**, LYN **(I)**, or YES **(L)** to β-actin in **(A, D, G, J)**.

Data in **(B-C, E-F, H-I, K-L, N)** were expressed as mean ± SD, and *p*-values were calculated via one-way ANOVA followed by Dunnett´s test. The individual replicate data in (**B**), (**C**), (**E**), (**F**), (**H**), (**I**), (**K**), and (**L**) are listed in **Table S10**. (ns, not significant; * $p < 0. 05$, ** $p < 0.01$)

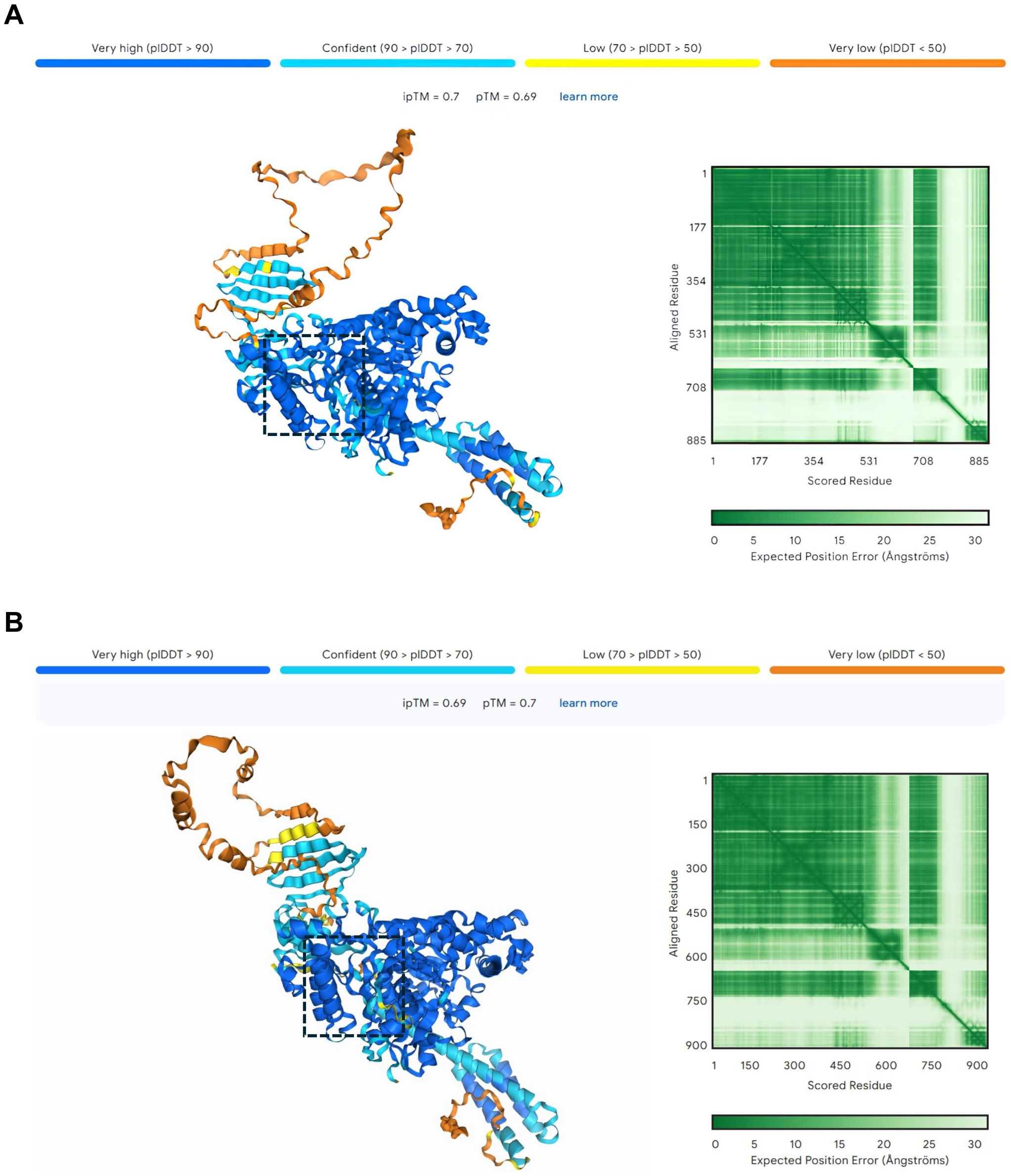


**Figure S6. Quality assessment of AlphaFold 3-predicted structures of the HSPA8-DNAJB6b complex before and after DNAJB6b Y53 phosphorylation, related to Figure 7.**
**(A)** Predicted structure and confidence metrics of the HSPA8-DNAJB6b complex in the unphosphorylated state. **(B)** Predicted structure and confidence metrics of the complex following DNAJB6b Y53 phosphorylation. Both models are color-coded by the predicted local distance difference test (pLDDT) scores, as indicated by the accompanying gradient scale. Heatmaps show the predicted aligned error (in Å), ranging from light green (higher error) to dark green (lower error). Black boxes highlight the region containing Y53 in **(A)** and pY53 in **(B)**, both of which were predicted with high confidence.

Fig. 1A

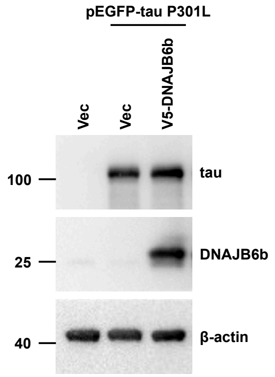


Fig. 1C

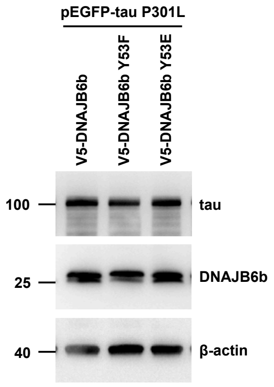


Fig. 1G

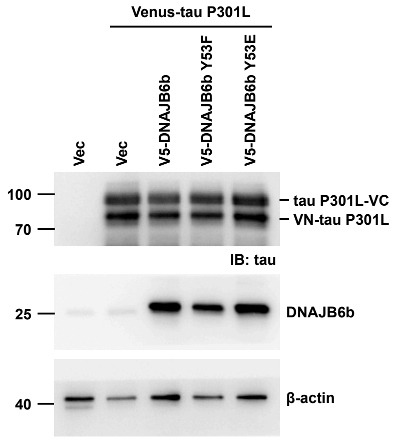


Fig. 2A

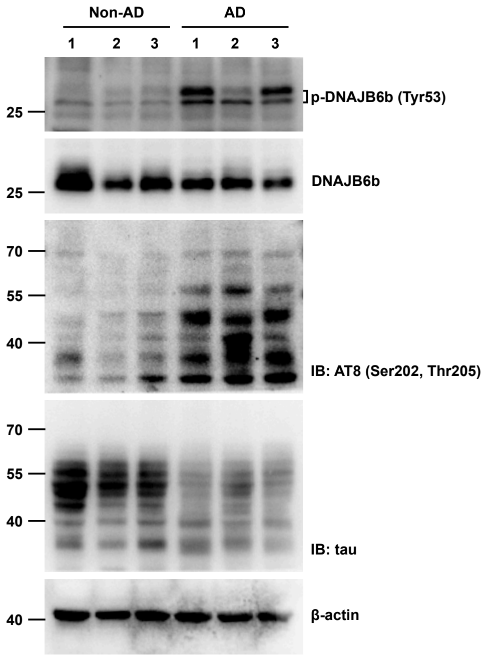


Fig. 3A

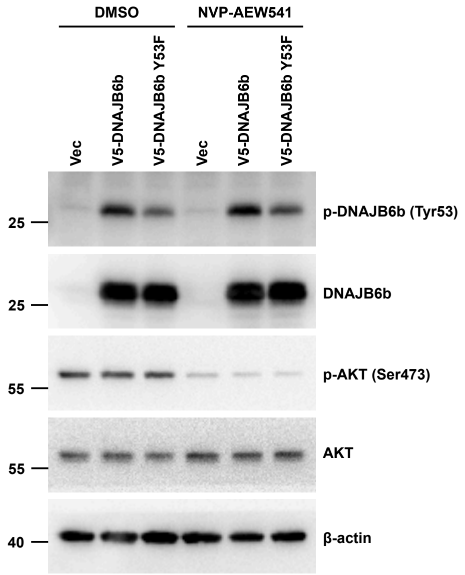

Fig. 3D

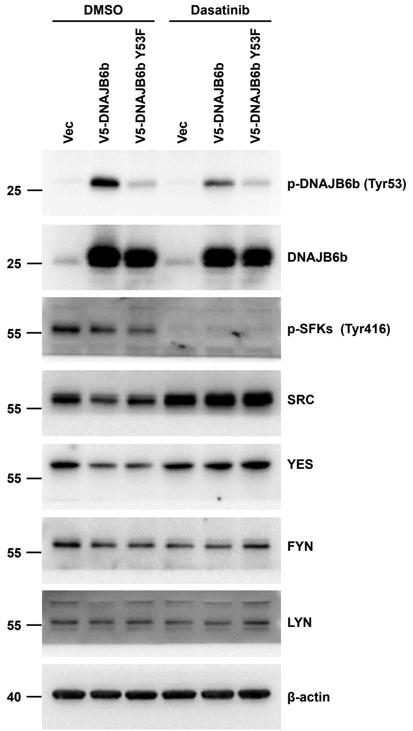


Fig. 4A

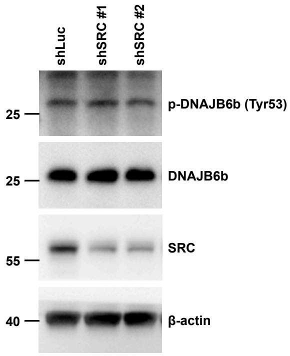


Fig. 4D

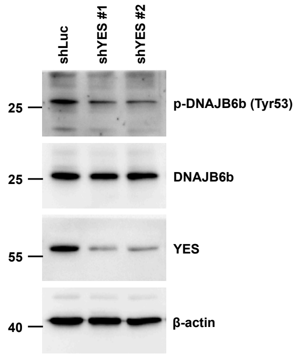


Fig. 4G

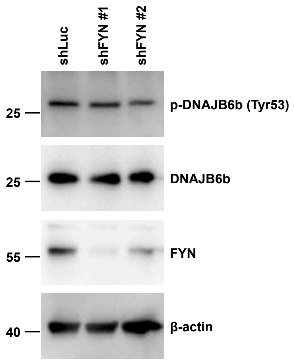


Fig. 4J

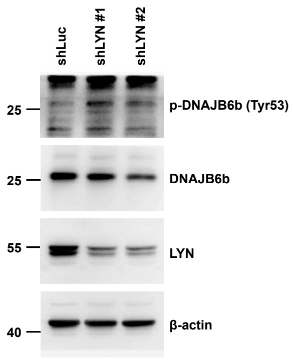


Fig. 4M

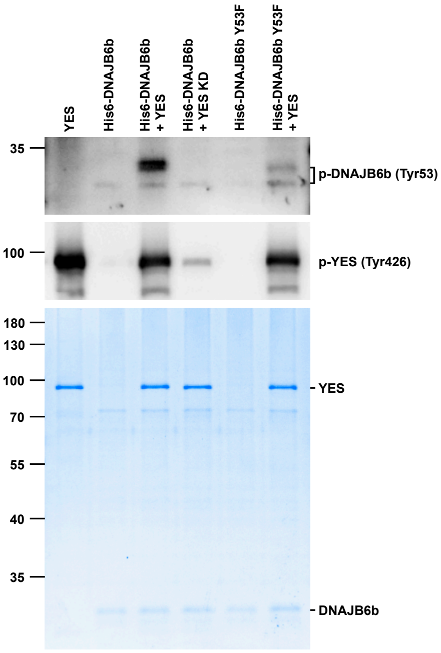


Fig. 5A

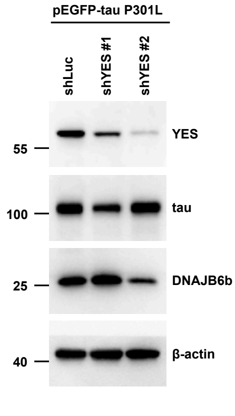


Fig. 5D

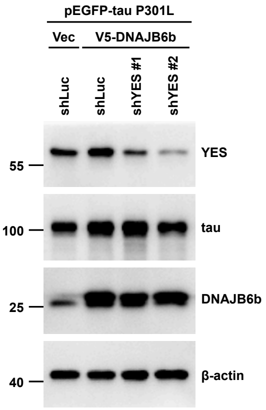

Fig. 7A

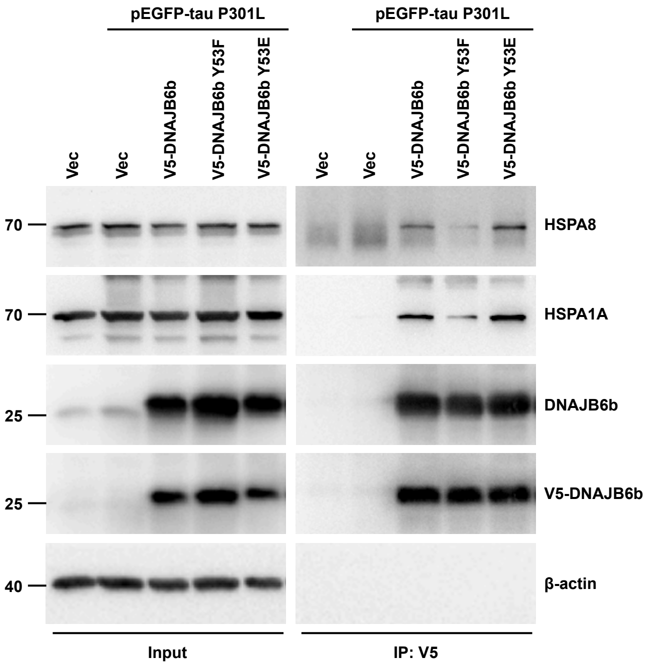


Fig. 7E

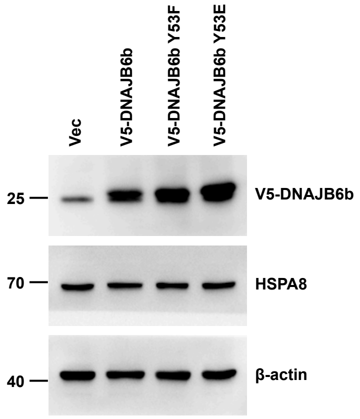


Fig. S1A

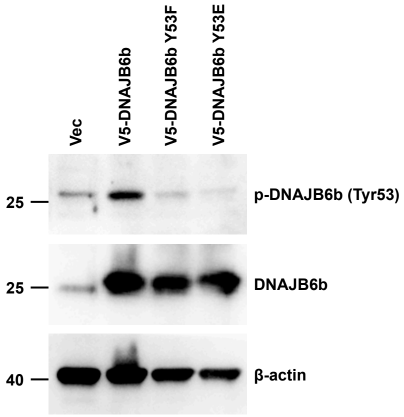


Fig. S2A

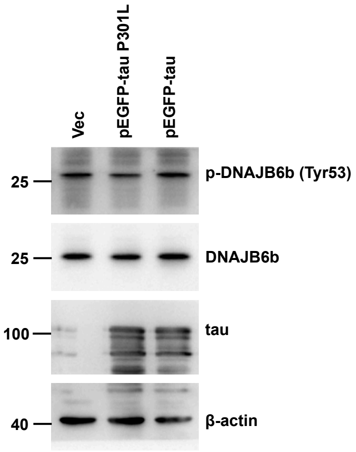

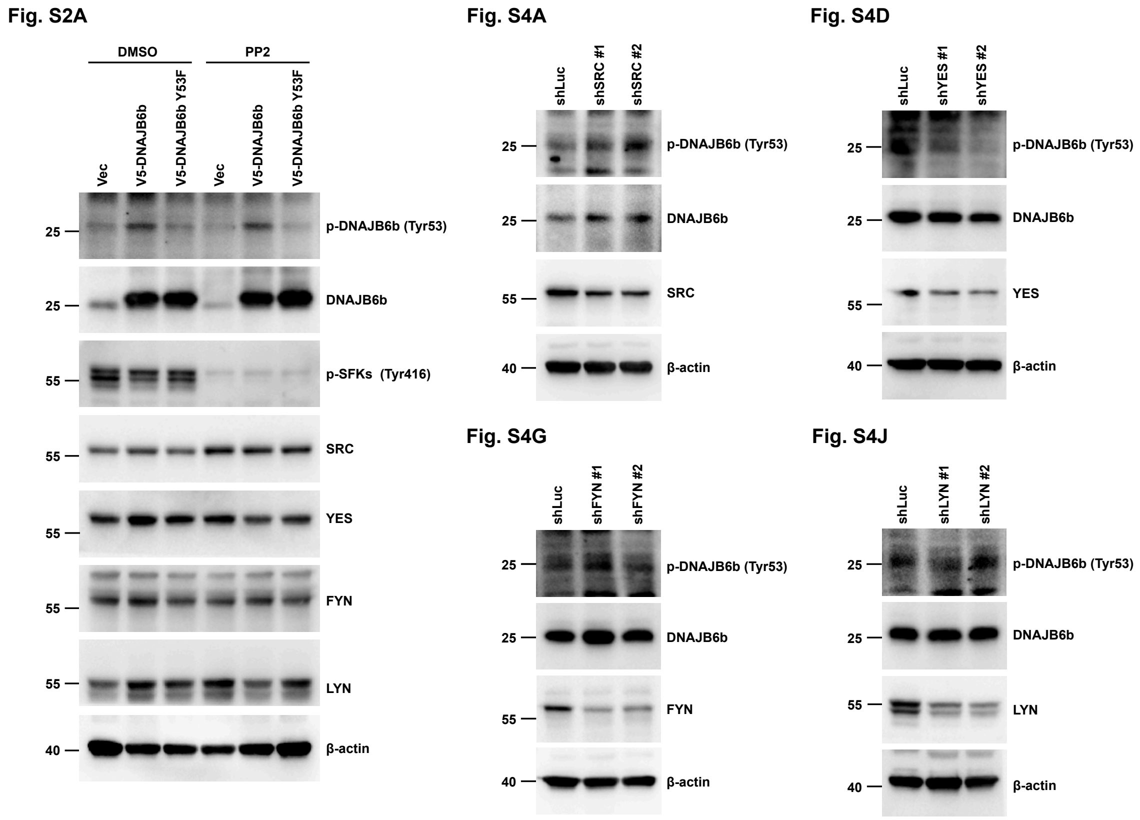


**Figure S7. Images of the full immunoblots.**

**Table S1. The phosphorylation sites on DNAJB6 were reported in the PhosphoSitePlus database.**

| Site | Amino acid sequence of DNAJB6 | LTP | HTP |
|---|---|---|---|
| Y4 | ____MVD(pY)YEVLGVQ | 0 | 1 |
| S15 | LGVQRHA(pS)PEDIKKA | 0 | 7 |
| Y53 | FKQVAEA(pY)EVLSDAk | 0 | 14 |
| S57 | AEAYEVL(pS)DAKKRDI | 0 | 1 |
| Y65 | DAKKRDI(pY)DKYGKEG | 0 | 4 |
| S226 | EEDGQLK(pS)LTINGVA | 0 | 1 |
| T228 | DGQLKSL(pT)INGVADD | 0 | 1 |
| S267 | PKPPRPA(pS)LLRHAPH | 0 | 3 |
| S277 | RHAPHCL(pS)EEEGEQD | 0 | 24 |
| S316 | KQKQREE(pS)KKKKSTK | 0 | 1 |

Candidates of the DNAJB6b PTM site are assessed through the available data from PhosphositePlus® (https://www.phosphosite.org/homeAction). Phosphorylation sites on DNAJB6b were listed in the chart. Low-throughput papers (LTP) indicate that the number of records in which this modification site was determined using methods other than discovery mass spectrometry. High-throughput papers (HTP), on the other hand, report the number of records in which this modification site was assigned solely by proteomic mass spectrometry.

**Table S2. Prediction of possible upstream kinases for DNAJB6b by NetPhos 3.1.**

| Phosphorylation site | Sequence (N→C) | Score | Predicted kinase |
|---|---|---|---|
| Y53 | VAEA**Y**EVLS | 0.555 | unsp |
| | | 0.412 | INSR |
| | | 0.408 | SRC |
| | | 0.358 | EGFR |

Possible upstream kinases responsible for the phosphorylation of DNAJB6b at Y53 are listed in the chart. The closer the probability score is to 1, the more likely it is that the residue is phosphorylated by the predicted kinase. In this case, candidates with scores higher than 0.4 are listed. INSR has a score of 0.412, and SRC has a score of 0.408 for phosphorylating DNAJB6b Y53. NetPhos 3.1 server is available at: http://www.cbs.dtu.dk/services/NetPhosK/. (unsp: non-specific prediction)

**Table S3. The primers used for cloning, related to STAR Methods.**

| Primers | Sequence | Source |
|---|---|---|
| DNAJB6-Y53F-BstBI-For | 5’GCAAGTAGCGGAGGCATTCGAAGTGCTGTCGGATGC3’ | IDT |
| DNAJB6b-Y53E-NdeI$^{del}$-For | 5’GCAAGTAGCGGAGGCAGAAGAAGTGCTGTCGGATG3’ | IDT |

**Table S4. The oligonucleotide sequences of shRNA, related to STAR Methods.**

| Oligonucleotide sequences of shRNA | Source | Identifier |
|---|---|---|
| pLKO.1-shLuc<br>Target seq.: 5’GCGGTTGCCAAGAGGTTCCAT3’ | Academia Sinica | TRCN0000072249 |
| pLKO.5-shLuc<br>Target seq.: 5’ GCGGTTGCCAAGAGGTTCCAT3’ | Academia Sinica | TRCN0000231719 |
| pLKO.1-shSRC #1<br>Target seq.: 5’ GCTGACAGTTTGTGGCATCTT3’ | Academia Sinica | TRCN0000199313 |
| pLKO.1-shSRC #2<br>Target seq.: 5’ CATCCTCAGGAACCAACAATT3’ | Academia Sinica | TRCN0000195339 |
| pLKO.1-shFYN #1<br>Target seq.: 5’ GCGATCAGCAAACATTCTAGT3’ | Academia Sinica | TRCN0000003097 |
| pLKO.1-shFYN #2<br>Target seq.: 5’ GTGCCAACAATCCTAGTGCTT3’ | Academia Sinica | TRCN0000003101 |
| pLKO.5-shLYN #1<br>Target seq.: 5’ GGAATCCTCCTATACGAAATT3’ | Academia Sinica | TRCN0000218210 |
| pLKO.5-shLYN #2<br>Target seq.: 5’ GAGTGACGATGGAGTAGATTT3’ | Academia Sinica | TRCN0000230901 |
| pLKO.1-shYES #1<br>Target seq.: 5’ CCAAAGTCAGAATTGCTCAAA3’ | Academia Sinica | TRCN0000121242 |
| pLKO.1-shYES #2<br>Target seq.: 5’ GCTGTCATTATTTCCTCTTAT3’ | Academia Sinica | TRCN0000121062 |

**Table S5. The plasmids used in this study**, **related to STAR Methods.**

| Plasmid | Source | Identifier |
|---|---|---|
| pEGFP-C1 | Clontech | Cat# 6084-1 |
| pEGFP-C1-tau P301L (2N4R) | (Chang et al., 2023) | N/A |
| VN-tau (P301L) | Addgene | Cat# 87634 |
| tau (P301L)-VC | Addgene | Cat# 87633 |
| pcDNA5/FRT/TO-V5 control | (Chang et al., 2023) | N/A |
| pcDNA5/FRT/TO-V5-DNAJB6b | Addgene | Cat#19528 |
| pcDNA5/FRT/TO-V5-DNAJB6b Y53F | This study | N/A |
| pcDNA5/FRT/TO-V5-DNAJB6b Y53E | This study | N/A |
| pET-28a-DNAJB6b | This study | N/A |
| pET-28a-DNAJB6b Y53F | This study | N/A |

**Table S6. The antibodies used in this study, related to STAR Methods.**

| Antibodies | Source | Identifier |
|---|---|---|
| DNAJB6 | Abcam | Cat# ab198995<br>RRID: AB_2924896 |
| HSPA8 | Novus | Cat# NB120-2788<br>RRID: AB_2120309 |
| HSPA1A (HSP70 C92F3A-5) | Santa Cruz Biotechnology | Cat# sc-66048<br>RRID: AB_832518 |
| tau | GeneTex | Cat# GTX112981<br>RRID: AB_10730753 |
| Phospho-Tau (Ser202, Thr205)<br>Monoclonal Antibody (AT8) | Thermo Fisher Scientific | Cat# MN1020<br>RRID: AB_223647 |
| V5 | Abcam | Cat# ab9116<br>RRID: AB_307024 |
| V5 | Cell Signaling Technology | Cat# 13202<br>RRID: AB_2687461 |
| V5 | Thermo Fisher Scientific | Cat# R960-25<br>RRID: AB_2556564 |
| β-actin | Proteintech | Cat# 60008-1-Ig<br>RRID: AB_2289225 |
| GFP | GeneTex | Cat# GTX113617<br>RRID: AB_1950371 |
| Phospho-AKT (Ser473) | Cell Signaling Technology | Cat# 9271<br>RRID: AB_329825 |
| AKT | Cell Signaling Technology | Cat# 9272<br>RRID: AB_329827 |
| Phospho-Src Family (Tyr416) | Cell Signaling Technology | Cat# 2101<br>RRID: AB_331697 |
| SRC | Cell Signaling Technology | Cat# 2108<br>RRID: AB_331137 |
| FYN | GeneTex | Cat# GTX109428<br>RRID: AB_10729465 |
| Phospho-LYN (Tyr397) | Abcam | Cat# ab300118<br>RRID: AB_3683527 |
| LYN | GeneTex | Cat# GTX101222<br>RRID: AB_10720516 |
| YES | Cell Signaling Technology | Cat# 3201<br>RRID: AB_11178531 |

| | | |
|---|---|---|
| Peroxidase AffiniPure® Goat Anti-Rabbit IgG (H+L) | Jackson ImmunoResearch | Cat# 111-035-003 RRID: AB_2313567 |
| Peroxidase AffiniPure® Goat Anti-Mouse IgG (H+L) | Jackson ImmunoResearch | Cat# 115-035-003 RRID: AB_10015289 |
| Alexa Fluor™ 594 Goat anti-Mouse IgG | Invitrogen | Cat# A-11005 RRID: AB_2534073 |

**Table S7. The human brain samples used in this study.**

| Catalog | Lot # | Sex | Age | Ethnicity | Pathology |
|---|---|---|---|---|---|
| **NB820-59177** | C511134 | Male | 26 | Asian | Normal clinical diagnosis |
| **NB820-59177** | C807591 | Male | 82 | Caucasian | Normal clinical diagnosis |
| **GTX28771** | 822400674 | Male | 82 | Caucasian | Aortic stenosis |
| **NB820-59363** | C511136 | Female | 93 | Hispanic | Alzheimer's disease, Lymphoma |
| **NB820-59363** | C511137 | Male | 87 | Caucasian | Alzheimer's disease, Prostate cancer |
| **GTX26622** | 822304853 | Female | 93 | N/A | Alzheimer's disease, Lymphoma |

120 **Table S8. The chemicals and reagents used in this study, related to STAR Methods.**

| Chemicals and Reagents | Source | Identifier |
|---|---|---|
| Dulbecco's Modified Eagle Medium/Nutrient Mixture F-12 (DMEM/F12) medium | Cytiva | Cat# SH30023.02 |
| Antibiotic/Antimycotic Solution (100x) | Capricorn Scientific | Cat# AAS-B |
| Avantor® Seradigm, Select Grade Fetal Bovine Serum (FBS) | VWR | Cat# 89510-186 |
| Puromycin | Sigma-Aldrich | Cat# P8833 |
| Dimethyl sulfoxide (DMSO) | Sigma-Aldrich | Cat# D8418-100ML |
| NVP-AEW541 | MedChemExpress | Cat# HY-50866 |
| Dasatinib | MedChemExpress | Cat# HY-10181 |
| PP2 | MedChemExpress | Cat# HY-13805 |
| beta-Amyloid Peptide (1-42) (human) | Abcam | Cat# ab120301 |
| Recombinant Human/Murine/Rat BDNF, PeproTech® | Thermo Fisher Scientific | Cat# 450-02-10UG |
| RIPA Lysis Buffer, 10X | Merck Millipore | Cat# 20-188 |
| cOmplete™ EDTA-free Protease Inhibitor Cocktail | Roche | Cat# 4693132001 |
| PhosSTOP™ inhibitor tablets | Roche | Cat# 4906837001 |
| Protein G Mag Sepharose Xtra magnetic beads | Cytiva | Cat# 28967070 |
| TALON® Superflow™ histidine-tagged protein purification resin | Cytiva | Cat# 28957499 |
| Amicon Ultra-0.5 Centrifugal Filter Unit (3 kDa) | Merck Millipore | Cat# UFC500396 |
| Human YES1, GST Tag Recombinant Protein | Thermo Fisher Scientific | Cat# A15559 |
| PageRuler™ Prestained Protein Ladder (10-180 kDa) | Thermo Fisher Scientific | Cat# 26616 |
| GXBio Prestained Protein Ladder (25-300 kDa) | Biohelix | Cat# XP25300 |
| Luminata™ Crescendo Western HRP Substrate | Merck Millipore | Cat# WBLUR0500 |
| Opti-MEM® I Reduced Serum Medium | Gibco | Cat# 31985070 |
| Lipofectamine™ LTX reagent with PLUS™ reagent | Thermo Fisher Scientific | Cat# 15338100 |

| | | |
|---|---|---|
| Bio-Rad Protein Assay | Bio-Rad Laboratories | Cat# 5000006 |
| Duolink® In Situ Wash Buffers, Fluorescence | Sigma-Aldrich | Cat# DUO82049 |
| Duolink® In Situ PLA® Probe Anti-Mouse PLUS | Sigma-Aldrich | Cat# DUO92001 |
| Duolink® In Situ PLA® Probe Anti-Rabbit MINUS | Sigma-Aldrich | Cat# DUO92005 |
| Duolink® In Situ Detection Reagents Red | Sigma-Aldrich | Cat# DUO92008 |
| DAPI | Thermo Fisher Scientific | Cat# D1306 |
| Fluoromount™ Aqueous Mounting Medium | Sigma-Aldrich | Cat# F4680 |
| Cover glasses circular, 15 mm Ø No. 1 (0.13-0.16 mm) | Marienfeld Laboratory Glassware | Cat# 0111550 |

121

**Table S9. The instruments and software used in this study, related to STAR Methods.**

| Instrument and software | Source | Identifier |
|---|---|---|
| ChemiDoc™ Imaging System | Bio-Rad | Cat# 12003153 |
| 96-well dot-blot apparatus | Bio-Rad | Cat# 170-6545 |
| ApoTome.2 microscope | Zeiss | N/A |
| Zen Blue 2.6 | Zeiss | N/A |
| ImageJ | NIH | https://imagej.net/ |
| Image Lab | Bio-Rad | https://www.bio-rad.com/en-tw/product/image-lab-software?ID=KRE6P5E8Z |
| BioRender | N/A | https://biorender.com/ |
